# Early-life human lymphopoiesis generates a distinct B-cell lineage

**DOI:** 10.64898/2026.10.07.756737

**Authors:** Alexandra F. Nazzari, Krishna Kothari, Junkai Yang, Astrid Kosters, Palak Chaudhry, Ian A. Glass, Birth Defects and Research Laboratory (BDRL), Eliver E. B. Ghosn

**Affiliations:** Department of Medicine, Division of Immunology, Lowance Center for Human Immunology, Emory University School of Medicine, Atlanta, GA 30322, USA; Emory Vaccine Center, Emory University School of Medicine, Atlanta, GA 30322, USA; Department of Pediatrics, Division of Rheumatology, Emory University School of Medicine, Atlanta, GA 30322, USA; Department of Pediatrics, University of Washington, Seattle, WA 98195, USA; Seattle Children’s Research Institute, Seattle, WA 98145, USA

## Abstract

We characterize the first wave of human B-lymphopoiesis across gestation (n=42 donors) and identify a developmentally and functionally distinct B-cell lineage that shapes antibody specificity before birth. The earliest lymphoid progenitors engage an evolutionarily conserved genetic program enriched for imprinted genes associated with embryogenesis in mice and humans. These progenitors arise in the yolk sac and acquire Early B-cell Factor 1 (EBF1) in the embryonic liver, initiating B-lymphopoiesis before Hematopoietic Stem Cells emerge. Functionally, these early-life B-lymphoid progenitors (eBLPs; CD81^hi^TdT^−^IL3R^+^) respond to IL-3 to promote B-cell fate commitment and generate two waves of stereotyped public B-cell receptor (BCR) repertoires, shaping adaptive immunity at birth. The same molecular program and public BCRs are recapitulated during B-cell differentiation from pluripotent stem cells in vitro and after transplantation into humanized mice, where they generate antibody-secreting cells and SOX4^+^ B cells resembling infant lymphocytes. Developmental origin therefore intrinsically programs B-cell identity and antibody specificity.

## Introduction

Human B-cell development begins early in gestation^1,2^, primarily in the fetal liver (fL)^3–5^, which is the main site of hematopoiesis during the first trimester, before the bone marrow niche is established^6–8^. Although fL-exclusive B-cell progenitors have been identified in humans (CD7^+^CD10**^−^**)^9–12^, their origins, developmental requirements, and the functional properties of their progeny remain unresolved. Understanding how human early-life B cells develop and function would shed light on how evolution has shaped the immune system to adapt to changing environmental pressures throughout the organism’s life and provide insights into how adaptive immunity operates during childhood.

In mice, B cells develop in sequential “layers” of functionally distinct lineages, with the first emerging in the fL and later in the adult bone marrow (aBM)^13^. B-cell progenitors in the fL harbor a fetal-exclusive genetic program^14–16^, marked by *Lin28b* expression, and give rise to B-1a. By contrast, the aBM mainly produces conventional B cells, known as follicular B-2. Unlike conventional B cells, fetal B-1a progenitors develop in the extra-embryonic yolk sac before the hematopoietic stem cell (HSC) emerges (HSC-independent)^13,17–29^, and generate public natural antibodies that recognize self (including apoptotic cells), commensals, and pathogens^30–34^, thereby maintaining homeostasis and serving as the first line of defense against infections. Since the murine B-1a lineage was originally identified by comparing B-lymphopoiesis in fL versus aBM^35^, we aimed to characterize the first wave of human B-cell development in fL (n=42 donors) during the first two trimesters of pregnancy and to compare its developmental program and progeny with those of the aBM. We generated multi-omics single-cell datasets and provide integrated fL and aBM maps as an interactive atlas that can be explored in our user-friendly web application, SuPERR^36^, to reveal mechanisms of B-cell development across the lifespan.

We show that the first surge of human B-lymphopoiesis during gestation mirrors that of mouse B-1a cells, including expression of key regulators LIN28B, ARID3A, and IGF2BP3, giving rise to a functionally distinct population of human B-1 cells before birth. Notably, the earliest lymphoid progenitors that emerge after conception co-opt an evolutionarily conserved imprinted gene program associated with embryogenesis in mice and humans. This “primitive” program marks lymphoid progenitors that first appear in the extra-embryonic yolk sac and the EBF1^+^ B-cell progenitors that colonize the embryonic liver before HSCs emerge. Unlike aBM, early-life B-lymphoid progenitors (eBLP; CD81^hi^TdT^−^IL3R*^+^*) respond to IL-3 and generate two sequential waves of stereotyped public BCR repertoires, characterized by unique CDR3 motifs produced by a cell-intrinsic molecular program that can be recapitulated in vitro and in vivo. When transplanted into humanized mice, fL progenitors generate the same public BCRs and differentiate into SOX4^+^ mature B cells that resemble infant naïve B cells. Thus, our studies reveal a developmentally and functionally distinct B-cell lineage that represents the human counterpart to mouse B-1a, linking developmental origin to B-cell identity and antibody specificity, and providing a framework for understanding how human B-1 shapes adaptive immunity and B-cell-mediated diseases across the human lifespan.

## Results

### Progression of B-lymphopoiesis across gestation

We first characterized human B-lymphopoiesis during active fL hematopoiesis^3^, 7 to 20 weeks post-conception (wpc) (n=42 donors) (Figure 1A). By 7 wpc, we observed the complete B-cell developmental pathway in the fL, from CD34^+^ progenitors to mature B cells (IgM^+^IgD^+^) (Figure 1B and Figures S1A-S1C). The total number of developing B cells (CD19^+^) increases rapidly from <5% at 7 wpc to over 20% of the total CD45^+^ immune cells in the fL by 10 wpc (Figure 1B). To characterize the gradual progression of B-cell development across gestation, we defined three stages: fL-I (7-12 wpc), fL-II (13-15 wpc), and fL-III (16-20 wpc). These timeframes capture the change in the ratio of CD34^+^ progenitors to IgM^+^IgD^+^ mature B cells (Figure 1B and Figure S1D). CD19^+^CD34^+^ B-progenitors are detected throughout the three stages (7-20 wpc), but their proportion declines over time (CD34^+^: 44% in fL-I to 11% in fL-III). Conversely, mature B cells constitute a small but readily detectable population in fL-I that gradually increases and stabilizes after 16 wpc (IgM^+^IgD^+^: 21±7%) (Figure S1D). Having defined the dynamics of fL B-lymphopoiesis, we selected ten fL samples spanning these gestational stages for in-depth molecular and functional characterization.

**Figure 1.**
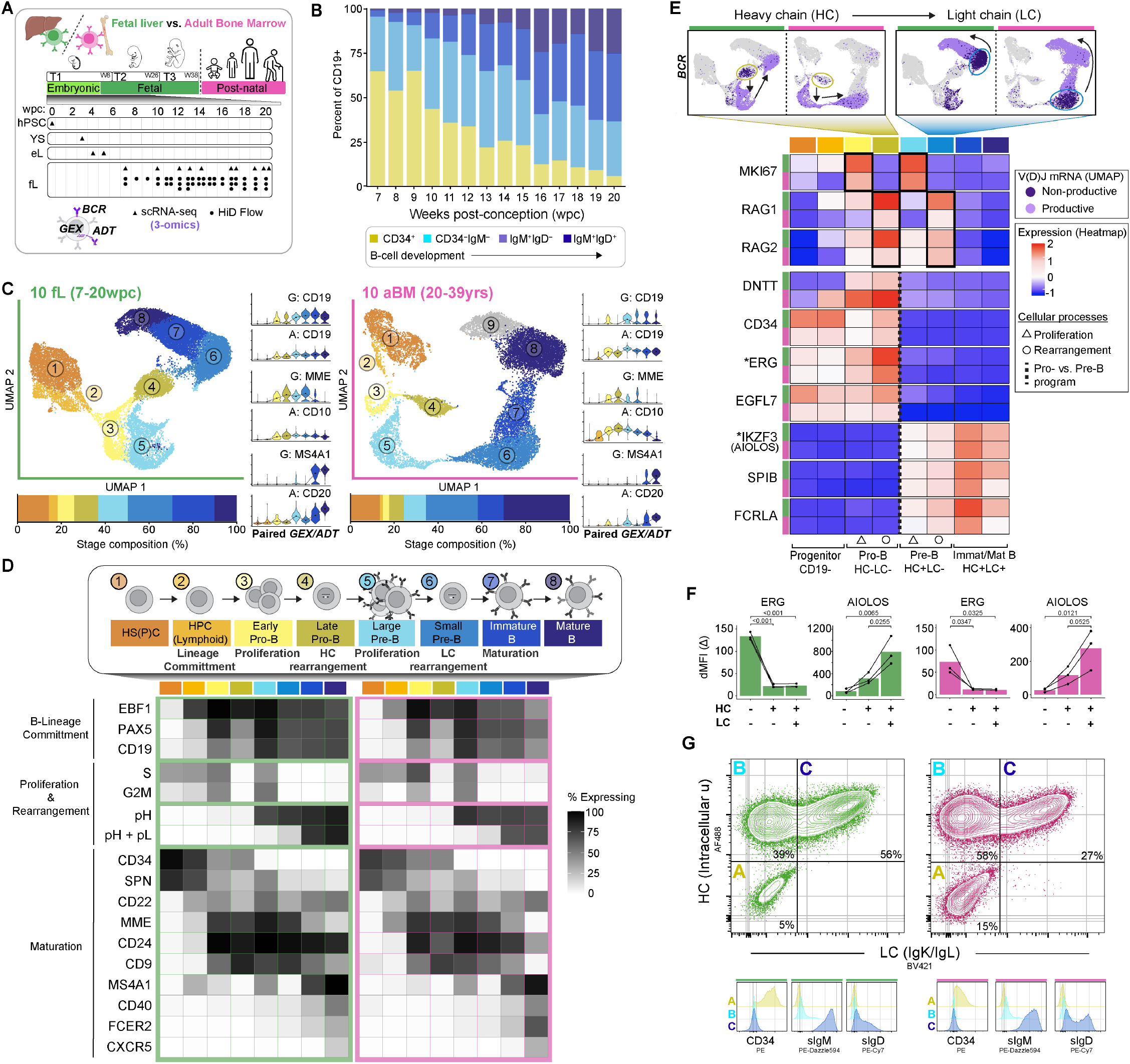
Multi-omics characterization of the B-cell developmental trajectory in early life versus adulthood. (A) Multi-omics characterization of human B-cell development across gestation using 3-omics: gene expression (GEX), cell-surface protein by Antibody-Derived Tags (ADT), and B-cell receptor (BCR) sequencing. (B) Flow cytometry characterization of fetal liver (fL) B-cell development (CD19+ population). Average composition of developmental subsets binned by weeks post-conception (wpc) (n=42 donors). (C) UMAP embeddings of hematopoietic progenitors and developing B cells in human fetal liver (left, n=10 donors, 24,298 cells) and adult bone marrow (aBM) (right, n=10 donors, 17,638 cells). Bar plot depicts the average cluster distribution; individual samples are shown in **Figure S1C.** The UMAP cluster colors are used in panels d-e (green: fL, magenta: aBM). Gene expression (G/GEX) and matching cell-surface protein by antibody-derived tags (A/ADT) from representative fL and aBM donors (right of corresponding UMAP). The black circles indicate median log-normalized GEX or ADT expression. (D) B-cell stage definitions exemplified by key biological processes. Heatmap shows the percentage of cells expressing each transcript displayed by UMAP cluster. BCR status determined by BCR-seq V(D)J matrix; productive heavy chain (pH) and light chain (pL). (E) Unproductive and productive BCR transcripts displayed on fL and aBM UMAPs. Heatmap depicts average gene expression (z-score, log-normalized expression scaled across both tissues). (F) Intracellular staining of ERG and Aiolos in CD19^+^ B cells (n=3 fL, n=3 aBM). Stages determined by HC expression (intracellular mu) and LC expression (surface kappa or lambda). Bars indicate the mean of the delta median fluorescence intensity (MFI) value compared to the fluorescence-minus-one (FMO) control. Paired measures within a sample are connected by a black line. Statistical test performed using a linear mixed-effects model with developmental stage as a fixed effect and donor as a random effect. Pairwise stage comparisons were performed using estimated marginal means with multiplicity-adjusted p-values (Tukey method). (G) Staging by BCR using flow cytometry in developing B cells (CD19^+^CD24^hi^CD38^+^). Representative sample shown (n=3 fL, n=3 aBM). Histograms display expression levels in the corresponding gated population.

### A core gene program conserved across the lifespan directs stepwise B-lymphopoiesis

To enable direct comparisons between fetal and adult B-lymphopoiesis, we generated multi-omics single-cell maps (total gene expression (GEX), cell-surface proteins by antibody-derived tags (ADT), and B-cell receptor repertoire (BCR-seq); 10X Genomics) of the human fL (n=10, 7-20 wpc, 24,298 cells) and the adult bone marrow (aBM) (n=10, 20-39 years old, 17,638 cells) (Figure 1C; Tables S1, S2). To increase B-cell recovery at early gestational ages and capture rare progenitor populations, we enriched hematopoietic stem and progenitor cells (HS(P)Cs) and B cells by immunomagnetic depletion of non-B-cell lineages (STEMCELL Technologies) before encapsulation. After integrating and clustering fL and aBM samples (Methods and Figures S2A, S2B), as described previously^36,37^, we identified eight matching clusters corresponding to key molecular events defining each stage of B-cell development (Figures 1C, 1D and Figures S2C-S2J; Table S3). We first identified the three core molecular programs of B-lymphopoiesis: I. lineage commitment, II. proliferation and BCR formation, and III. maturation. Lineage commitment is driven by master transcription factors, notably Early B-cell Factor 1 (EBF1), which marks the earliest B-cell specification program within the hematopoietic progenitor cells (HPCs, cluster #2), followed by Paired Box 5 (PAX5) and CD19^38,39^ (Figure 1D). After lineage commitment, a highly regulated process with alternating steps of cell proliferation and RAG-dependent V(D)J gene rearrangement forms the BCR, which determines antibody specificity. To establish the developmental trajectory, we relied on well-characterized molecular events that track progressive V(D)J gene recombination^40,41^. The progression from heavy chain (HC) V-D-J recombination in pro-B to light chain (LC) V-J recombination in pre-B defines BCR formation (Figures 1D, 1E). More specifically, we tracked HC and LC gene rearrangements from *non-productive* to *productive* V(D)J mRNA transcripts by BCR-seq to infer progressive B-cell developmental stages (Figure 1E and Figure S2E), a method that we^36^ and others^42^ previously described as a reliable, biologically driven approach to order the B-cell developmental trajectory.

By integrating multi-omics datasets (BCR, GEX, ADT), we identified stark transcriptional reprogramming at the pro-B to pre-B transition, coinciding with expression of a productive HC (Figure 1E and Figure S2F). We identified 46 and 51 genes, respectively, that distinguish pro-B (e.g., DNTT, CD34, EGFL7, SOCS2, CD99, ERG^43^, ELK3^44^) from pre-B cells (e.g., IKZF3 (Aiolos)^45^, SPIB^46^, FCRLA^47^, TCL1A^48^, IGLC1), in both fL and aBM (Table S4). We validated the pro-B and pre-B gene programs at the protein level by intracellular staining of ERG and Aiolos, respectively (Figure 1F and Figure S3A). Following productive rearrangement of both HC and LC, maturation proceeds, as revealed by gradual changes in B-cell maturation genes (e.g., SPN, MME, MS4A1, CD40, FCER2) and corresponding cell-surface proteins (Figure S2C). Thus, using stage-wise multi-omics analysis, we identified conserved gene sets that define the B-cell developmental pathway in both fL and aBM (Figures 1D, 1E and Figure S2D; Table S3).

Despite similar expression of core gene programs between fL and aBM, we identified key differences that distinguish fetal B-lymphopoiesis. In contrast to aBM, the earliest B-progenitor in fL lacks DNTT, the gene encoding terminal deoxynucleotidyl transferase (TdT) (Figure 1E and Figure S2G). TdT generates diversity in BCR sequences by catalyzing non-templated nucleotide additions (N-additions) between the V-(D)-J genes. Although DNTT is absent in the earliest fL B-progenitor (cluster #2), its expression begins in the subsequent pro-B stages (clusters #3 and #4) but never reaches the high transcript levels observed in aBM (Figure S2G). Despite delayed DNTT expression, fL B-progenitors express higher levels of the recombinase genes RAG1 and RAG2 than aBM (Figure 1E), indicating a robust and perhaps premature start of HC recombination. Similarly, LC recombination initiates earlier in the fL trajectory (Figure 1E), prior to the expected small pre-B stage (cluster #6). Specifically, LC transcripts are detected in the fL as early as large pre-B (cluster #5) and are marked by the cyclin gene CDKN1C (Figure S2H), a fetal-specific imprinted gene that negatively regulates cell proliferation^49^. This premature LC recombination and distinct cell-cycle regulation suggest a non-canonical pre-BCR program in fetal B cells, similar to what has been described for mouse fetal B-1a^50^.

During LC recombination, we identified IGF2 as the key gene marking the transition from kappa to lambda LC gene usage, which may include secondary rearrangements and receptor editing^51,52^, in small pre-B cells in both fL and aBM (Figure S2I). Notably, IGF2 is an imprinted gene involved in fetal growth^53^ and is expressed more broadly in fL than in aBM (Figure S2I), likely explaining the increased lambda LC usage in early-life B cells^54^ (Figure S2J). Finally, after productive HC and LC rearrangement, fL B cells express an IgM^hi^IgD^lo^ cell-surface phenotype during the transition from immature to mature B cells, a stage marked by alternative splicing from IgM to IgD (Figure 1G and Figures S3B, S3C), suggesting differences in BCR regulation similar to those observed in murine fetal B-1a cells^20,55^.

We provide here the integrated fL and aBM maps as an interactive atlas that can be further explored to reveal mechanisms of B-cell development across the lifespan. The data can be explored within SuPERR^36^, our web-based R Shiny application for multi-omics scRNA-seq analysis and visualization (instructions in Figures S4A-S4C).

### Evolutionarily conserved imprinted gene program shapes early-life B-lymphopoiesis

In mice, the first wave of fL B-lymphopoiesis is regulated by a distinct gene program that gives rise to a developmentally and functionally distinct B-cell lineage, B-1a^30,56,57^. To investigate whether the gene program regulating human fL B-lymphopoiesis mirrors that of fetal B-1a development, we performed stage-wise differential gene expression (DGE) analysis comparing fL and aBM (Figure 2A). We identified 317 genes uniquely expressed in fL and 158 genes uniquely expressed in aBM B-lymphopoiesis (Figure S5A; Table S5). The genes upregulated during fL B-lymphopoiesis were associated with three main programs: I. epigenetics and transcription, II. cell cycle and translation, and III. cell signaling and BCR modulation (Figure 2A). Notably, fetal B cells express unique cell-cycle genes located within imprinted loci clusters^58^ that regulate embryogenesis^49,59^ (IGF2, H19, KCNQ1OT1, and CDKN1C on chromosome 11; DLK1 and MEG3 on chromosome 14) (Figure 2A), suggesting that early-life immune cell development co-opted imprinted genes involved in organogenesis and fetal growth^53,60^. The genes involved in cell signaling include regulators of BCR lipid raft organization (ARID3A^61^, EFHD2^62^, MARCKS^63^, and CD81^64^ and phosphoinositide 3-kinase (PI3K) (PDE3B^65^), as well as phosphatases (DUSP2^66^, PTPRK^67^, and PTPRJ^68^), which together may regulate BCR signaling thresholds in human fL B cells, similar to mouse B-1a^16,68–70^. The fetal-exclusive gene program in human B cells also includes expression of key regulators of mouse B-1a progenitors, namely LIN28B, ARID3A, and IGF2BP3^14–16^, present in human fL progenitors across gestation (Figure 2B). To further identify genes conserved between human and mouse fL B-lymphopoiesis, we analyzed scRNA-seq datasets^71^ of mouse HPCs from fL (embryonic day 16.5) and aBM (8 weeks old) (Figure 2C; Table S6). Strikingly, we found a conserved lymphoid gene program between the two species, including orthologous imprinted genes (H19, Cdkn1c, Dlk1, Meg3) (Figure 2C) involved in fetal growth, suggesting evolutionarily conserved mechanisms of embryogenesis and hematopoiesis in both mice and humans.

**Figure 2.**
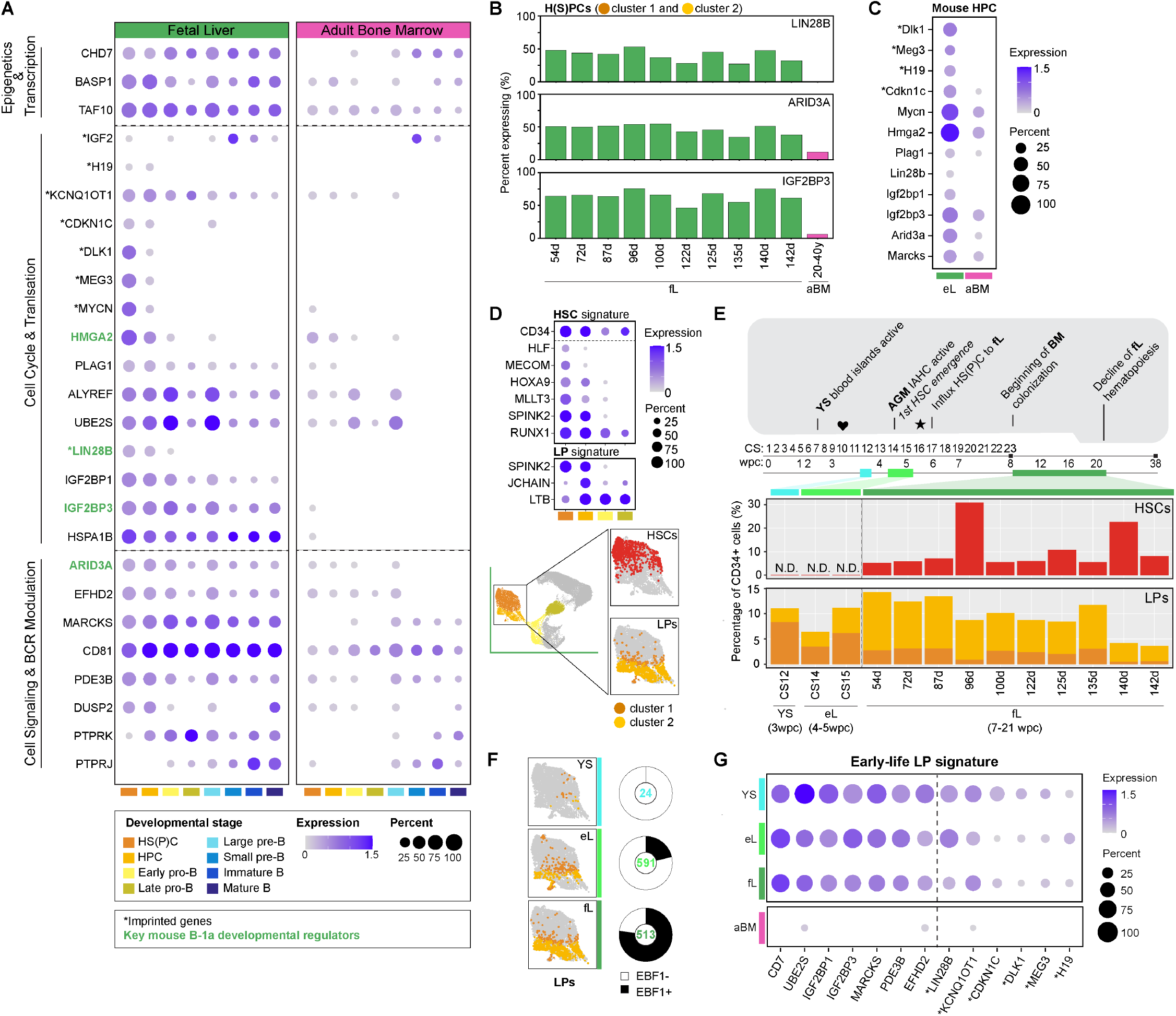
Evolutionarily conserved lymphoid progenitors engage genes associated with embryogenesis and initiate B-lymphopoiesis before HSCs emerge. (A) Fetal-exclusive genes identified by differential gene expression analysis. (B) Expression of essential mouse B-1a progenitor genes in human hematopoietic progenitors across gestational age. (C) Select fetal-exclusive genes highly expressed in hematopoietic progenitor cells (HPCs, Lineage^−^Sca1^+^c-kit^+^CD48^+^) from mouse embryonic liver (eL, E16.5) compared with mouse adult bone marrow (aBM, 8 weeks)^71^. Imprinted genes in mice are denoted with an asterisk^58^. (D) Lymphoid progenitor (LP: LTB^+^SPINK2^+^JCHAIN^+^) and hematopoietic stem cell (HSC: RUNX1^+^HOXA9^+^MLLT3^+^ MECOM^+^HLF^+^SPINK2^+^) identified by gene expression and highlighted on the fL UMAP. (E) Quantification of HSCs and LPs across gestational age. Gestational timeline indicates key developmental events, including initiation of heartbeat (♥) and establishment of definitive fetal circulation (★)^72,74,75^. (F) Early-life LP signature across tissues and age. Yolk sac (YS, CS12), embryonic liver (eL, CS14 and CS15), and fetal liver (fL, 7-20wpc) samples were mapped to the fetal liver UMAP from Figure 1C. The number of progenitors identified per tissue is labeled inside the pie, with the percentage of cells expressing the EBF1 transcript shaded black. (G) Dot plot displays the top differentially expressed genes in LPs from early life compared with adult BM (aBM LP, n=73. Early-life LP cell count displayed in Figure 2f). Imprinted genes in humans are denoted with an asterisk^58^.

### The first wave of B-lymphopoiesis initiates before HSCs

While the extra-embryonic yolk sac (YS) is the first site of blood cell development, the definitive HSC first emerges in the embryonic aorta-gonad-mesonephros (AGM) around 4 wpc (∼32 days post-conception)^72–74^. Subsequently, as the vascular system forms^75^, progenitors migrate and colonize the embryonic liver (eL) by 6 wpc^4,76^. To identify the first site of human lymphopoiesis, we analyzed publicly available datasets from the YS (Carnegie stage 12 (CS12), 3 wpc)^77^ and the embryonic liver (eL) (CS14 and CS15, 4-5 wpc)^76^, before the emergence of the first HSC. As expected, we did not detect definitive HSCs (HLF^+^ MECOM^+^ MLLT3^+^ HOXA9^+^ RUNX1^+^ SPINK2^+^)^76^ in the YS and eL (Figures 2D, 2E). In contrast, we found a robust population of lymphoid progenitors (SPINK2^+^LTB^+^JCHAIN^+^)^76^ as early as 3-5 wpc in the YS and eL (Figures 2D, 2E), suggesting HSC-independent lymphopoiesis in humans. To determine whether these YS and eL lymphoid progenitors express a gene program consistent with B-cell lineage specification, we further characterized them and compared them to our fL B-lymphopoiesis map (Figures 2F, 2G). Strikingly, the lymphoid progenitors in the YS, eL, and fL share a conserved gene program (CD7, UBE2S, IGF2BP1, IGF2BP3, MARCKS, PDE3B, and EFHD2) and engage the same imprinted genes associated with embryogenesis that define fetal B-lymphopoiesis in mice and humans (LIN28B, KCNQ1OT1, CDKN1C, DLK1, MEG3, and H19)^58^ (Figure 2G and Figure S5B; Table S7). These lymphoid progenitors express the B-lineage specification gene EBF1 as early as ∼4 wpc in the eL (Figure 2F), before HSCs appear. These results suggest an evolutionarily conserved imprinted gene program for generating early-life B cells, via a developmental pathway that initiates in the YS before definitive HSCs emerge.

### CD81^hi^TdT^−^IL-3R^+^ marks the earliest B-cell fate specification

We used nascent EBF1 and PAX5 expression to mark the onset of B-cell fate specification (Figure 3A and Figure S6A) and to compare EBF1^+^ early-life versus EBF1^+^ adult B-lymphoid progenitors (eBLP vs. aBLP). We identified eBLP as CD7^+^DNTT^−^MME^−^, contrasting with the aBLP CD7^−^DNTT^+^MME^+^ phenotype (Figures 3B, 3C; Table S8)^10,78^. Unlike mouse fetal B-1a progenitors, which lack DNTT^79^, human eBLP upregulates DNTT at the pro-B stage (Figure 3C and Figure S2G), suggesting differences in the perinatal BCR repertoire between humans and mice. Other genes associated with BCR regulation also distinguish eBLP, including expression of MARCKS^63^, ARID3A^61^ (Figure 3B), and high transcription of CD81^64^ (Figures S6A, S6B). In addition to BCR-associated genes, we identified IL3R as the primary cytokine receptor that marks eBLP (Figure S6C). Consistent with this, eBLP co-expresses the gene encoding the adapter protein downstream of IL-3R known as the cytokine-dependent hematopoietic cell linker (CLNK) (Figures 3B, 3C), suggesting active IL-3 signaling^80^. Thus, we defined human fL eBLP at the transcriptional level as CD7^+^DNTT^−^MME^−^CD81^hi^IL3R^+^.

**Figure 3.**
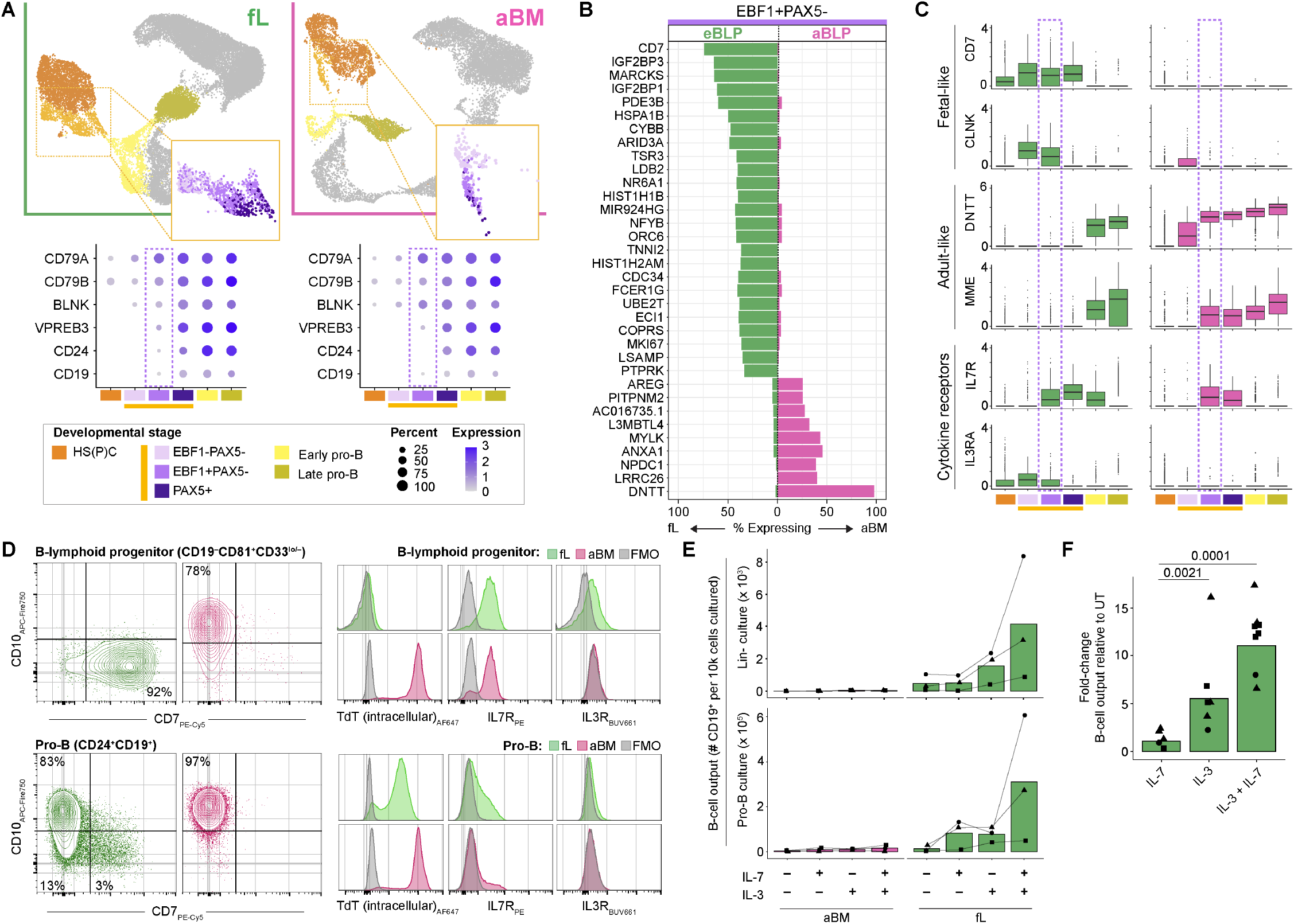
IL-3 requirement distinguishes early-life (eBLP) from adult (aBLP) B-lymphoid progenitors. (A) Subpopulations of hematopoietic progenitors (UMAP cluster 2) highlighted by EBF1 and PAX5 expression. Dot plot shows log-normalized transcript expression of B-cell-related genes that mark B-cell commitment in both fL and aBM. (B) Differentially expressed genes that distinguish early-life (eBLP) from adult (aBLP) B-lymphoid progenitors within the EBF1^+^PAX5^−^population (fL: 543 cells, aBM: 90 cells). (C) Boxplot displaying fetal and adult progenitor features. Log-normalized gene expression with center line marking median and whiskers extending to 1.5x interquartile range. EBF1, PAX5 subsets colored according to panel Figure 3A. Purple dashed box highlights eBLP (left, green) and aBLP (right, magenta). (D) Flow cytometry plots of CD34^+^ progenitor subsets, B-lymphoid (top) and pro-B cells (bottom) in fL (green) and aBM (magenta) distinguished by CD7 and CD10 expression (representative sample, n=3 fL, n=3 aBM donors). CD34^+^ progenitors were isolated by bead enrichment (STEMCELL Technologies) and gated as CD45^+^CD45RA^+^CD38^+^ cells, excluding lineage-specific markers (CD3^−^CD14^−^CD16^−^CD56^−^CLEC4C^−^CD19^−^). Pro-B cells were identified by CD24 and CD19 co-expression. B-lymphoid progenitors are contained within the CD19^−^CD81^hi^CD33l^o/-^ population. Flow cytometry histograms (right) show TdT, IL7R, and IL3R expression in fetal (green) and adult (magenta) B-lymphoid progenitor and pro-B cells. Fluorescence-minus-one (FMO) controls are shown in grey. (E-F) Lineage-negative (Lin^−^) progenitors and pro-B cells sorted by flow cytometry and cultured for 15 days with 10ng/ul of the respective cytokines (IL-3, IL-7). 3 independent experiments were performed (denoted by shape) with 1 to 4 replicates per condition per donor (n=3 fL, n=3 aBM donors). (E) Generation of CD19^+^ B cells from cultured progenitors. (F) Fold-change in CD19^+^ B-cell output from fL Lin^−^progenitors compared to untreated. Statistical test performed using a linear mixed-effects model with cytokine treatment as a fixed effect and donor as a random effect. Pairwise comparisons were performed using estimated marginal means with multiplicity-adjusted p-values (Tukey method).

Next, we characterized the eBLP phenotype by flow cytometry. Within the human CD45^+^CD34^+^ HS(P)Cs (CD3^−^CD14^−^CD16^−^CD56^−^CLEC4C^−^CD19^−^), we further distinguished the CD45RA^+^CD38^+^ compartment, which contains lympho-myeloid progenitors^81^ (Figure S6D). Because CD10 can distinguish lymphoid from granulocyte-monocyte progenitors in adult but not in fetal progenitors^10^, we tested whether CD81, which we found is highly expressed on EBF1^+^ cells (Figures S6E, S6F), could be detected on the cell surface and used to mark lymphoid progenitors^82^. Indeed, cell-surface CD81 protein segregates myeloid (CD81^lo/-^CD33^hi^) and lymphoid (CD81^hi^CD33^−^) compartments in both the fL and aBM (Figure S6D-S6F). Within the CD81^hi^ lymphoid compartment, we identify eBLPs by cell-surface CD7 expression and lack of CD10 (CD7^+^CD10^−^)^9–11^, whereas aBLPs are CD7^−^CD10^+^. Downstream of eBLPs, but before pro-B, we identified a fetal-exclusive intermediate stage, pre-pro-B (CD34^+^CD19^+^CD10^−^)^10,11^, that downregulates CD7 as it progresses toward the B-cell developmental trajectory (Figure 3D). We then validated DNTT expression by intracellular flow cytometry (i.e., TdT protein). Unlike in adult aBLP, TdT is absent in early-life eBLP but is upregulated at the pro-B stage, albeit at ∼5-fold lower levels than in aBM pro-B (Figure 3D, Figure S6G). Finally, consistent with gene expression, cell-surface IL-3R protein distinguishes eBLP from aBLP (Figures 3C, 3D). Interestingly, unlike in aBM, cell-surface IL-7R persists in fL from eBLP through the pre-pro-B and pro-B stages, suggesting distinct cytokine signaling dynamics. Thus, by flow cytometry, eBLP is contained within the CD7^+^TdT^−^CD10^−^CD81^hi^IL-3R^+^ compartment. We revised the current classification of human B-cell development to incorporate these findings and provide a diagram illustrating each stage, including cell-surface markers and gene expression profiles that distinguish fetal and adult B-lymphopoiesis (Figures S7A, S7B).

### IL-3 supports early-life B-cell lineage commitment

Because IL-3R and the downstream protein CLNK are expressed in eBLP but not in aBLP, we investigated the role of IL-3 in early-life B-lineage commitment and differentiation in vitro. We purified CD34^+^ HS(P)Cs and downstream pro-B cells (CD34^+^CD19^+^CD24^+^) by FACS to assess the role of IL-3 in B-lineage commitment and differentiation, respectively. These progenitors were purified from both fL and aBM and cultured for 15 days in StemSpan II serum-free media (STEMCELL Technologies) supplemented with IL-3 and/or IL-7 (Figures S8A, S8B). Strikingly, IL-3 substantially increased CD19^+^ B-cell production from HS(P)Cs in fL but not in aBM (Figure 3E). While IL-7 alone had limited effect on B-lineage commitment, adding it with IL-3 increased the number of CD19^+^ B cells by >10-fold in fL cultures (Figure 3F). As expected^83^, IL-3 also supported myeloid cell expansion in both fL and aBM progenitors (Figure S8C). At the pro-B stage, both IL-3 and IL-7 independently supported the survival, expansion, and differentiation of fL pro-B cells (Figure 3E). However, the strongest effect was observed when the two cytokines were combined, revealing synergy between IL-3 and IL-7 on pro-B cells (Figure 3E). Although flow cytometry detected cell-surface IL-3R only at the earlier eBLP stage, these findings suggest that IL-3 continues to influence fL B-lymphopoiesis at the downstream pro-B cells, potentially through low levels or rapid turnover of cell-surface IL-3R that was undetectable in our assays.

Given the effects of IL-3 and IL-7 on fL B-lymphopoiesis, we analyzed gene expression in our datasets and publicly available data^5^ to identify potential sources of these cytokines in the fL and aBM milieus. Although IL-7 is known to be produced by non-immune (CD45^−^) stromal cells^84^, we also detected IL-7 expression in B cells at the late maturation stage in the aBM, and to a lesser extent in fL (Figure S8D). In contrast, IL-3 transcripts in the fL were detected primarily in granulocytes, including mast cells, basophils, and eosinophils (Figure S8E). These findings reveal potential interactions between lymphoid and granuloid cells in the fL that promote B-cell development in early life. Overall, these results highlight IL-3 as a critical cytokine for early-life but not adult B-lymphopoiesis.

### Unlike aBM, the fL gene program promotes stereotyped public BCRs

We previously showed that mouse B-1a, but not B-2, generates a conserved public BCR repertoire defined by the heavy chain’s complementarity-determining region 3 (CDR3) amino acid (AA) sequences^34^. The CDR3 is the most diverse region of the BCR, formed by the junction of the V-(D)-J segments, and is critical for antibody specificity^85^. Because TdT, which contributes to CDR3 diversity, exhibits distinct expression in fL eBLP compared to pro-B cells, we characterized the BCR CDR3 AA sequences (HC and LC) across gestation. Strikingly, as in mouse B-1a, human B cells in fL, but not aBM, generate a public HC CDR3 (CDRH3) AA sequence (Figures 4A, 4B). Among B cells with a productive HC, we identified 96 public CDRH3 AA sequences (0.56% of total productive HC in fL) expressed by two or more fL donors (Figure 4B), including the ARDSSSWYYFDY motif identified in 5/10 fL (but not adult) donors. These fL public antibody CDRH3s persist through B-cell maturation, with 39 of the public sequences expressing productive HC and LC (Figure S9A). In contrast to the CDRH3, which generates public BCR only across fL donors, the LC CDR3 (CDRL3) is broadly shared across both fL and aBM donors (Figure 4B and Figure S9B).

**Figure 4.**
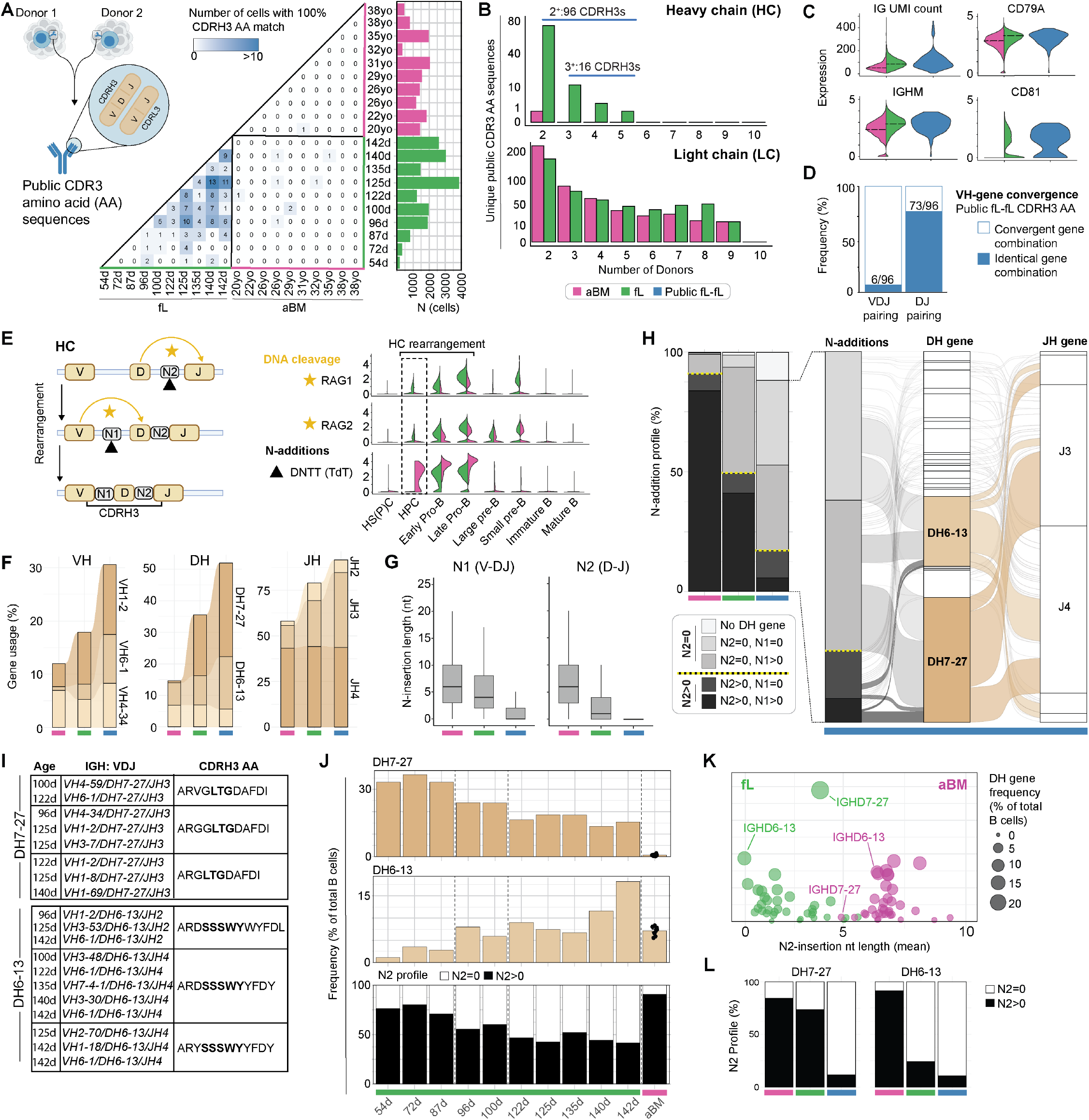
Early-life gene program generates two waves of stereotyped public BCRs before birth. (A) Pairwise comparison of exact CDRH3 amino acid (AA) sequence matches between donors, defined as public BCR (blue). For CDRH3 analyses, we included all B cells with a productive heavy chain (HC) (n=17,377 fL and n=12,998 aBM cells). Class-switched B cells were excluded to analyze recently formed B cells. (B) Number of unique CDRH3 (top) and CDRL3 (bottom) AA sequence matches found between two or more donors. For CDRL3 analysis, we included all cells with both productive HC and light chain (LC) transcripts (9,879 fL and 11,125 aBM cells). (C) BCR components expressed in B cells with productive HC and LC from aBM (magenta) and fL (green) and public fL (blue). IG UMI count represents the total UMI count of productive HC and LC contigs. IGHM, CD79A, and CD81 display gene expression (log-normalized). (D) VDJ-gene convergence is defined by the different VDJ-gene family combinations that result in the same AA sequence. 6/96 CDRH3s display identical V-D-J pairing, with the remaining formed by V-gene convergence. 73 of 96 public CDRH3 motifs are defined by the same DH-JH gene combinations. (E) VDJ rearrangement and CDRH3 schematic, including non-templated nucleotide (N)-additions (N1 and N2) (left panel). Gene expression (log-normalized transcript) levels of key enzymes essential for V(D)J rearrangement (RAG1, RAG2, DNTT; right panel) in fetal (green) and adult (magenta) B-lymphopoiesis according to stages in Figure 1C. Dotted box highlights the absence of DNTT on fetal hematopoietic progenitor cells (HPC). (F) HC gene family usage frequency. The top VDJ families expressed in the public fL-fL repertoire are shown. (G) N1- and N2-insertion length (number of nucleotides, nt) in cells with productive HC. The boxplot shows the interquartile range, with the median. (H) Bar plot (left) shows N-addition prolife in aBM (magenta), fL (green), and public fL (blue) cells. The dotted yellow line indicates the split between cells with or without N2-additions. A subset of productive HC denoted “No DH gene” is productive but lacks identifiable nucleotides between the annotated VH and JH segments. The alluvial plot (right) expands on the properties of the fL public CDRH3s (blue). Each line represents one cell and connects the N-addition profile to the corresponding DH and JH gene usage. The most frequently used D-genes (DH7-27 and DH6-13) are highlighted in tan. (I) Select public CDRH3 AA sequences from the fL, exemplifying DH7-27 and DH6-13 motifs (bold letters). (J) Frequency across gestation: DH7-27 gene usage, DH6-13 gene usage, and N2 insertions in the total B-cell population of each donor. The aBM values represent the average of all ten samples. (K) Mean N2-insertion nt length with respect to DH gene usage. (L) N2-addition profile within the DH7-27 and DH6-13 expressing cells in aBM (magenta), fL (green), and public fL (blue).

To further validate the extent to which early-life B cells generate a public BCR repertoire, we explored several publicly available BCR sequences from human prenatal, neonatal, and infant B cells. Strikingly, we found matching CDRH3 AA motifs in all early-life B-cell datasets analyzed^86–88^. Furthermore, we identified human CDRH3 AA motifs that overlap with those in mouse fetal B-1a cells (including ARRDY, ARLDY, ARGNY, ARGDY)^34^. Notably, the public CDRH3s are biochemically distinct, displaying a high Boman index (Figure S9C), indicative of increased protein interaction potential^89^, which likely contributes to the polyreactivity observed in prenatal antibodies^54^. The B cells expressing a public BCR also show high levels of immunoglobulin transcripts per cell and of the BCR-associated components CD79A and CD81 (Figure 4C).

We explored potential molecular mechanisms underlying public BCR generation, including V-D-J gene usage and N-additions at the V-to-D (N1) and D-to-J (N2) junctions (Figures 4D–4G and Figure S9D, S9E). The public BCR repertoire was largely formed through VH-gene convergence, in which different VH gene segments generate the same CDRH3 AA motifs (Figure 4D). Similarly, in mouse fetal B-1a, over 90% of public CDRH3s arise through VH-gene convergence^34^. Of the 96 human public CDRH3s, we identified only six clones with identical V-D-J gene combinations across all donors: VH1-2/DH7-27/JH4, VH6-1/DH6-13/JH4, VH1-69/DH6-6/JH1, VH3-23/--/JH3, VH3-53/DH7-27/JH3, and VH4-34/DH7-27/JH3. Over 75% of public BCRs lack D-to-J N2 additions (N2=0), reflecting the absence of TdT expression at the eBLP stage, when D-to-J joining likely initiates. In contrast, only <15% of total aBM B cells lack N2 additions (Figures 4G, 4H). Although fL public BCRs lack N2 additions, they still exhibit V-to-D (N1) additions, likely due to the upregulation of TdT in fL pro-B cells. As a result, ∼50% of human public CDRH3s display N-additions largely driven by N1 (Figure 4H).

### Public BCR repertoire distinguishes two sequential waves of B cells before birth

We observed two waves of biased usage of proximal V-D-J gene segments, most notable in the DH gene families closest to the JH segments, with DH7-27 and DH6-13 accounting for 46% of public BCRs (Figure 4H), leading to stereotyped BCR sequences (Figure 4I). The first wave of B-cell development in early gestation (7-12 wpc) is dominated by DH7-27 (Figure 4J), a fetal-biased gene^90^ that encodes an AA motif (LTG/LG) conserved across species, including the orthologous DQ52 in mouse fetal B-1a^91^. Surprisingly, although DH7-27^+^ B cells have the highest rate of N2 (D-to-J) additions^90^ (72% N2>0) (Figures 4K, 4L), the public BCRs expressing DH7-27 largely lack N2 (N2=0) (Figure 4L). Moreover, DH7-27 preferentially pairs with the VH1-2 gene. VH1-2, like DH7-27, is over-represented in early gestation and is expressed on >25% of all B cells formed at ∼7 wpc, resulting in over-representation of the VH1-2/DH7-27 pair in public BCRs (Figure S9F).

The second wave of fL B-lymphopoiesis is marked by DH6-13, which increases later in gestation (>12 wpc) and coincides with decreased DH7-27 usage (Figure 4J). Unlike DH7-27, DH6-13^+^ B cells show the lowest N2 insertion profiles during gestation (76% N2=0) (Figures 4K, 4L). This profile is exclusive to fL B cells, as >90% of aBM B cells expressing DH6-13 contain N2 insertions (Figure 4K). Therefore, the distinct BCR repertoire architecture in early versus late gestation defines two sequential waves of public BCRs marked by biased VH/DH gene usage and N1/N2 addition profiles. Notably, the bias we observed in VH and DH gene usage across gestation is not due to donor haplotype differences, as fL donors either carry the same gene allele or different alleles not expected to influence the observed pattern in VH and DH expression^92^ (VH1-2*02 detected in 9/10 fL donors; VH6-1*01, DH7-27*01, and DH6-13*01 alleles were detected in all fetal liver donors) (Figure S9G).

### A cell-intrinsic gene program determines B-cell identity and BCR repertoire

To determine whether the early-life B-cell program depends on the human fetal liver environment, we recapitulated B-lymphopoiesis in vitro and in vivo in humanized mice. In our previous study, we described a protocol for generating functional B cells and plasma cells in vitro from human pluripotent stem cells (hPSCs)^93^. Here, we performed single-cell multi-omics characterization of in vitro hPSC-derived B-lymphopoiesis and mapped these cells onto our fL and aBM atlases (Figure 1C), confirming their canonical developmental trajectory (Figures 5A, 5B, and Figure S10A). In vitro hPSC-derived B cells recapitulated the fL transcriptional program but not the aBM program, including expression of fetal-exclusive genes BASP1, ALYREF, IGF2BP3 and MARCKS (Figure 5C and Figure S10B). Strikingly, in vitro hPSC-derived B cells recapitulated the same fL public BCR architecture, including 66 identical CDRH3 AA sequences shared between hPSC- and fL-B cells (Figure 5D), suggesting that public BCR is not solely driven by antigen recognition within the fetal environment. Interestingly, the hPSC-derived B cells closely recapitulated the public BCR of early gestation (∼7-12 wpc), including a high frequency of DH7-27 and a low frequency of DH6-13 (Figures 5E–5H and Figures S10C, S10D). Accordingly, their N-addition profile matched that of DH7-27 from early gestation (N2>0) (Figures 5E, 5F). Thus, in vitro hPSC-derived B cells support a cell-intrinsic mechanism for the early-life B-cell program and provide further evidence of an early gestational wave of B-cell development that differs from that of late gestation.

**Figure 5.**
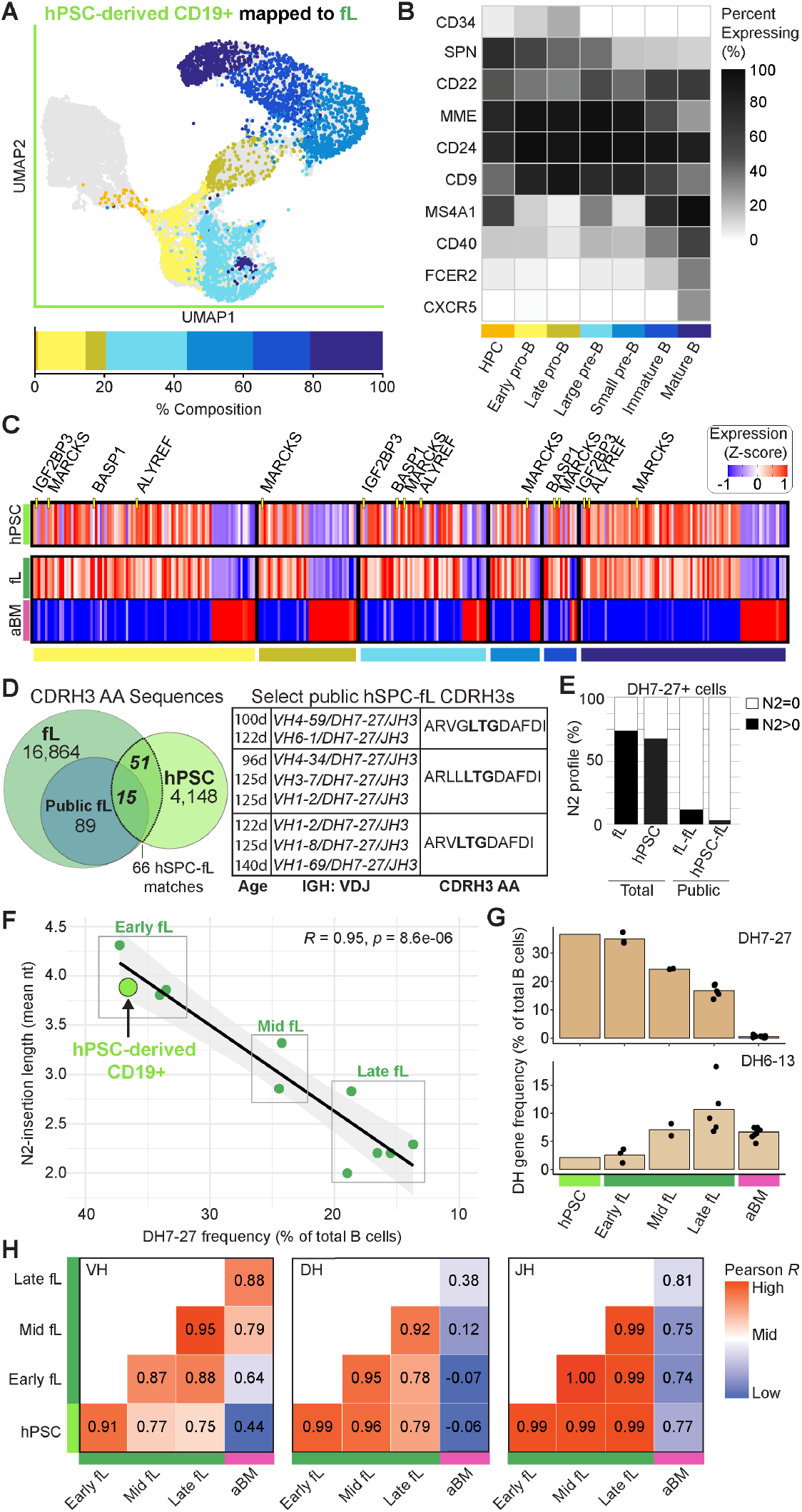
Cell-intrinsic fetal B-cell identity and public BCR recapitulated in vitro using human pluripotent stem cells (hPSC). (A) hPSC-derived B cells were mapped to the fL UMAP (grey) using Symphony integration (n=3,855 cells, GEX). Bar plot depicts cluster composition. Stages colored according to Figure 1C. (B) Heatmap shows the percentage of cells expressing B-cell maturation genes across development. (C) Heatmap representing the normalized gene expression (z-scored) in hPSC (light green), fL (green), and aBM (magenta) by developmental stages (clusters) colored according to the UMAP. Each column displays a gene from the differentially expressed genes list comparing fL and aBM. Select fetal-exclusive genes are labeled. (D) Exact CDRH3 amino acid (AA) sequences match between hPSC-derived B cells and fL. Exemplary DH7-27^+^ public CDRH3 AA sequences (DH7-27 motif in bold) listed. All hPSC-derived B cells with productive heavy chain (n=3,962 cells) were analyzed in panels **d-h**. (E) N2-addition profile within the total and public hPSC-derived B-cell pool expressing DH7-27. (F) Correlation of DH7-27 usage and mean N2 additions places hPSC-derived B cells with the early fL characteristics. (G) Frequency of DH7-27 and DH6-13 gene usage. (H) Pearson correlation for pairwise comparisons of heavy chain VH, DH, and JH gene usage. fL was segregated into early (7-12wpc: 54d, 72d, 87d), mid (13-15wpc: 96d, 100d) and late (16-20wpc, 122d, 125d, 130d, 135d, 142d) gestation according to Figure 1B.

Next, we transplanted 1.5 million fL CD34^+^ HS(P)Cs (142d donor) per mouse (n=9) (Figure 6A) and monitored CD19^+^ B-cell maturation in blood, spleen, and BM for over a year post-transplant (Figures 6B, 6C and Figures S11A-S11D). Notably, we found naturally occurring plasma cells/antibody-secreting cells (ASCs) (CD38^hi^CD27^+^) predominantly in the spleen, not the BM (Figure 6C). Class-switched circulating IgG and IgA antibodies (Figure 6B and Figure S11C) increased after ∼6 months and persisted for over a year. We next performed single-cell multi-omics characterization of splenic B cells and ASCs at 4 months post-transplant. As with hPSC-derived B cells, de novo B-lymphopoiesis in the humanized mice retained the fetal-exclusive gene program (Figure 6). We first identified 44 genes highly expressed in mature B cells in fL but not in aBM (Figure 6E, Table S9) and used them to generate a fetal score (UCELL; see Methods) to characterize B cells in the humanized mice (Figure S11E). De novo human B-lymphopoiesis in the mouse expressed the same fetal-exclusive genes identified in human fL (Figure 2 and Figure 6F), including UBALD2, TAF10, ALYREF, IGFBP3, MARCKS, DUSP2, and PTPRJ. Notably, we also identified SOX4 as a marker of fL-exclusive mature B cells (Figure S11F), and this pattern was recapitulated in the humanized mice (Figure 6F and Figure S11G). Interestingly, SOX4 was recently identified as a marker that distinguishes infant from adult naïve T cells^94,95^. We show that SOX4 also marks a subset of infant naïve B cells (Figure 6G and Figure S11H) that is enriched for a fetal transcriptional program (Figure 6G). Similarly, naturally occurring ASCs in humanized mice (Figure 6H and Figure S11I) were biased toward the fetal-exclusive program (Figure 6I), likely representing a source of natural antibodies similar to those in mouse B-1a^33,96^. Altogether, the in vivo findings further support a cell-intrinsic mechanism for the early-life B-cell program, which stably maintains a fetal-derived B-cell identity resembling that of infant naïve B cells and spontaneously differentiates into long-lived ASCs without exogenous stimuli.

**Figure 6.**
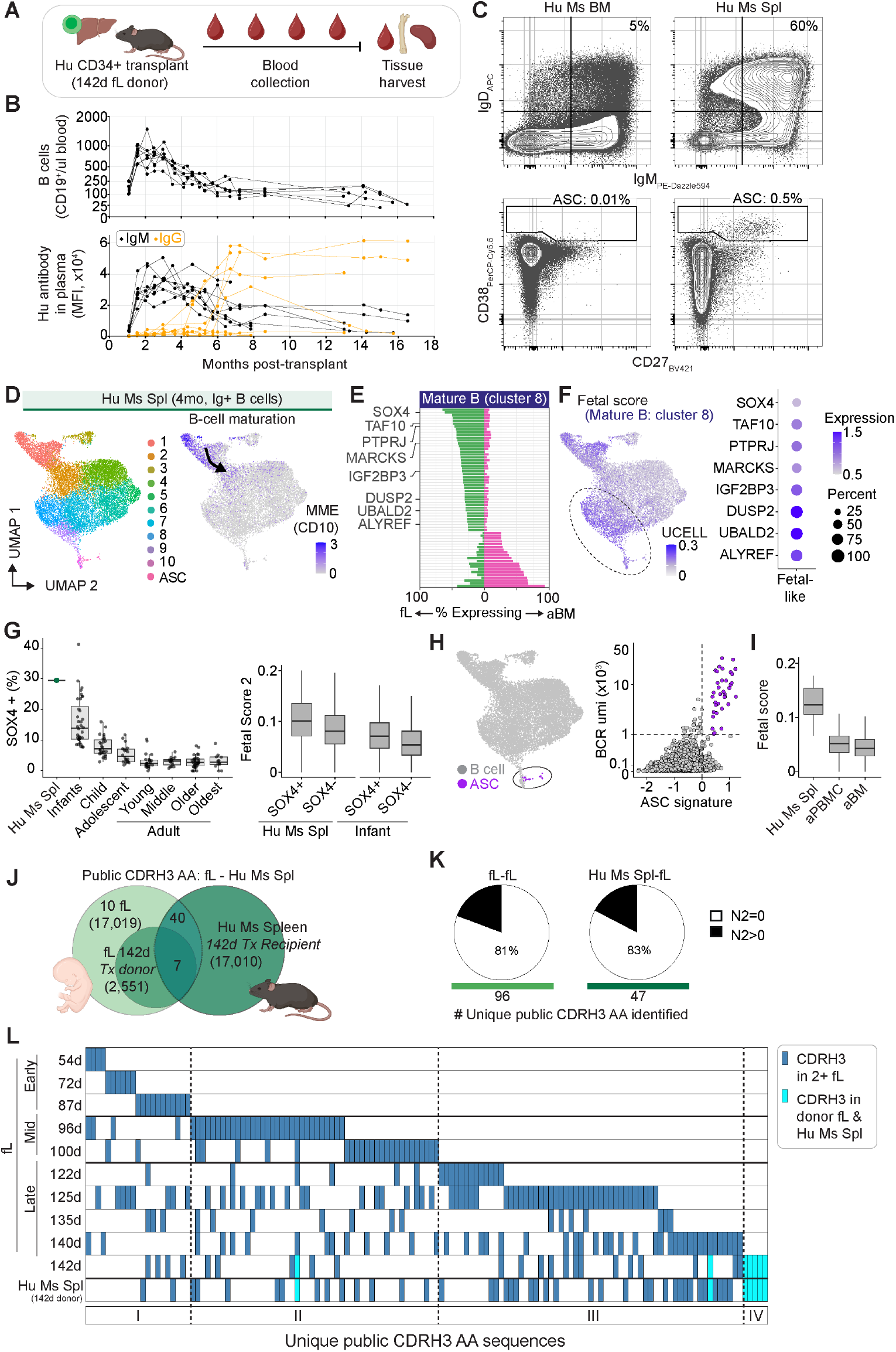
Adoptive transfers of fL CD34^+^ progenitors recapitulate fetal B-cell identity and generate the same stereotyped public BCRs. (A) Schematic for humanized mouse generation. fL CD34^+^ cells (1.5 million) were transplanted into triple-knockout mice (CD47^-/-^, RAG2^-/-^, IL2R gamma chain^-/-^) mice (n=9). (B) Absolute counts of CD19^+^ B cells in blood (top) and antibody (bottom), including IgM (black) and IgG (orange), in plasma assessed over 18 months post-transplant. Each line depicts an individual mouse (n=9). Antibody values are displayed as MFI with background subtraction using plasma from non-transplanted mice as controls (n=4 non-transplanted mice). (C) Flow cytometric characterization of human B cells and antibody-secreting cells (ASC) in mouse bone marrow (Hu Ms BM) and spleen (Hu Ms Spl) at four months post-transplant. (D) B cells from humanized mouse spleen at four months post-transplant (n=14,679 cells). UMAP displays clusters of mature B cells (Ig^+^ B cells denote productive heavy and light chains). MME (CD10) expression (log-normalized transcript) depicts B-cell maturation. (E) Differentially expressed genes in human fL and aBM mature B cells (cluster 8 in Figure 1). Gene selection is shown in **Table S9**. (F) Fetal score (UCELL, derived from the fL cluster 8 gene list) in B cells from a humanized mouse spleen. Clusters in the mouse spleen (panel D) with the highest fetal score are annotated as fetal-like (clusters 8-10). Selected fL genes expressed in fetal-like B cells are shown (full list in **Figure S11G**). (G) SOX4 expression in humanized mouse B cells (total mature cells, clusters 2-10) and human naïve B cells (0-105 years, n=167; infants: 2months-2years, child: 2-12 years, adolescent: 12-18 years, young adult: 18-40 years, middle-aged adult: 45-65 years, older adult: 65-85 years, oldest adult: 85-105 years). UCELL fetal score 2 was calculated for SOX4^+^ and SOX4^−^B cell subsets (we excluded SOX4 from the fetal score 2 gene list). (H) ASCs classified based on gene score signature (i.e., upregulation of XBP1, CD38, CD27, PIM2, SDC1, PRDM1 and downregulation of B-cell genes SPIB, BACH2, PAX5, FCER2, MARCH1, BANK1) and on total BCR/Ig UMI count (sum of total heavy and light chain transcripts). (I) Fetal score (UCELL) in ASCs derived from the Hu Ms Spl (n=36 ASCs), human adult (a)PBMC (3 donors, n=70 ASCs), and human aBM (10 donors, n=548 ASCs). (J) CDRH3 amino acid (AA) sequences of human B cells recovered from the Hu Ms Spl (dark green) compared to the 142d fL donor (green) and the total 10 fL donors (light green). 100% CDRH3 AA match is considered a public BCR. (K) N2 profile in public CDRH3 sequences. (L) Public CDRH3 AA sequences (blue) ordered by donor age and Hu Ms SPL. Each blue box represents a unique CDRH3 AA sequence in two or more individuals.

To further validate the cell-intrinsic mechanisms underlying public BCR generation, we characterized the BCR sequences of de novo human B cells engrafted in the humanized mouse. Remarkably, de novo human B-lymphopoiesis in mice recapitulated the same public BCR repertoire observed in the primary fL sample from the same donor (Figures 6J–6L). When we expanded the analysis to all 10 fL donors, we identified even more public BCRs, with most sequences lacking N2 insertions (N2=0), mirroring public BCRs in human fL (Figure 6K). Overall, although environmental factors play important roles in cell fate decisions, our findings challenge the notion that the human fetal environment alone shapes fetal B-lymphopoiesis. Instead, they point to an evolutionarily conserved cell-intrinsic mechanism that produces early-life public antibodies.

## Discussion

The newborn immune system is often considered functionally immature. However, recent studies in mice have challenged this notion, showing that immune cells rather emerge in sequential “layers” of functionally distinct populations adapted to each developmental age^13,19,25,97^. Here, we show that this “immune layering” also occurs in human B cells, with the first layer emerging in the embryonic liver before the first hematopoietic stem cell (HSC) develops. The first layer of human B-cell development mirrors that of mouse B-1a and operates through an evolutionarily conserved program driven by fetal-exclusive imprinted gene loci^49,58–60^ (parental allele silencing of LIN28B, IGF2, H19, DLK1, MEG3, KCNQ1OT1, CDKN1C, among others) involved in organogenesis, suggesting that early-life hematopoiesis co-opted gene programs that regulate embryonic growth. This fetal-exclusive program regulates B-cell responses in ways that distinguish them from their adult counterparts, including genes we identified involved in cell signaling and survival, and a cell-intrinsic program that gives rise to public antibodies, collectively shaping adaptive immunity and antibody specificity at birth.

Whether the fetal B cells we identified here migrate to specific tissues and persist to participate in B-cell-mediated immune responses into adulthood is difficult to investigate in humans. However, their cell-intrinsic program is enriched for genes involved in proliferation, high metabolic activity, and survival (IGF2BP3, MYCN, ALYREF, LIN28B), suggesting they may persist into postnatal life. Indeed, we show that fL CD34^+^ progenitors generate fetal-derived B cells that persist for over a year in humanized mice, where they maintain their early-life gene signature and spontaneously differentiate into antibody-secreting cells. Notably, the fetal-derived mature B cells in humanized mice express a gene signature recently described for infant naïve T cells, including SOX4^94,95^, suggesting a conserved early-life program that shapes the infant’s adaptive immunity, including B and T cell function. Another line of evidence for the postnatal persistence of fetal-derived B cells comes from leukemias. Certain aggressive forms of B-cell chronic lymphocytic leukemia (B-CLL) express fetal-exclusive stereotyped BCR clonotypes that are absent from the adult bone marrow^87,98^. B-cell acute lymphoblastic leukemia (B-ALL) also shares a molecular signature of the fetal-exclusive B-cell program we described here, including IGF2BP3^99^, MYCN^100^, ALYREF^101^, LIN28B^102^, and a subset of B-ALL is characterized by the expression of IL-3R^103,104^.

While IL-7 is often considered essential for B-cell development, we identified IL-3 as a key cytokine that promotes B-cell commitment in early life but not in adult B-cell progenitors. IL-3R marks the earliest fL CD34^+^ hematopoietic progenitors that express the B-cell specification gene EBF1, which we used to characterize the emergence of B-lymphoid progenitors (BLP) in early life (eBLP) and in adults (aBLP). Although eBLP (IL-3R^+^) and aBLP (IL-3R^−^) differ in IL-3R expression, they show similar IL-7R levels, consistent with the IL-7R^+^ multi-lymphoid progenitor (MLP)^81^ and early lymphoid progenitor (ELP)^1,9,10^ that display strong biases toward B-cell fate commitment. The differential expression of CD7, CD10, and TdT that we observed in eBLP versus aBLP is consistent with the previously described ontogenic “switch” that occurs within the IL-7R^+^ ELP^1,9–11,81^. Thus, the BLPs we characterize represent a defined subset within the previously described IL-7R^+^ ELP and are defined here by nascent EBF1 expression. In addition, fL B-lymphopoiesis downstream of eBLP gives rise to an intermediate population that precedes pro-B cells, named pre-pro-B (CD34^+^CD19^+^CD10^−^)^10,11^, which is absent in the aBM. Altogether, we define the fetal B-lymphopoiesis trajectory as MLP→eBLP→Pre-pro-B→Pro-B, which is distinguished from the adult BM MLP→aBLP→Pro-B (see our revised human B-lymphopoiesis diagram with corresponding markers and nomenclature in Figure S7).

The seemingly deterministic, stereotyped public BCRs generated by fL B-lymphopoiesis may reflect evolutionarily conserved mechanisms underlying the layered immune system. While previous studies revealed biased V(D)J clonotypes in fetal B cells,^105–107^ we extended these findings by showing a cell-intrinsic molecular mechanism that promotes two sequential waves of public BCRs (CDRH3 motifs) driven by VH-gene convergence, shaping antibody specificity before birth. In mouse B-1a cells, these convergent CDRH3 motifs are positively selected by self and commensal antigens, often containing lipid and polysaccharide residues^30,34,108,109^. Positive selection by self-antigens in the fetal environment could partly explain the mechanisms underlying human public BCRs; however, our data showing that the fetal B-cell program and stereotyped public BCRs are recapitulated in vitro and in vivo challenge this assumption. We showed that identical CDRH3 motifs and N-additions can be generated de novo in vitro from purified hPSC cultures and by transplanting fL CD34^+^ HS(P)Cs into humanized mice. The first and second waves of public BCRs were also separately recapitulated in vitro and in vivo, suggesting that developmental origins and cell-intrinsic mechanisms, rather than environmental factors alone, can determine B-cell identity and its antibody specificities.

While our studies could not determine whether HSC-dependent “adult-type” B-lymphopoiesis also initiates in the fL, we did not detect the adult BLP gene signature in our datasets throughout gestation. Recent studies suggest that fL HSCs contribute minimally to fetal hematopoiesis^110,111^. Instead, they describe embryonic multipotent progenitors (eMPPs) that give rise to most immune cells in the fetus, while fL HSCs expand and later seed the BM niche for hematopoiesis after birth. Consistent with this, our data show that fL HSCs are largely engaged in a maturation and expansion program, suggesting that fetal B cells likely originate exclusively from the early-life B-cell program (eBLPs) we characterized here. If so, future studies should determine when and how the transition from fetal to adult hematopoiesis occurs, and which immune cell lineages are generated as the organism ages.

Applying an adult-centric immunological lens to immune function overlooks the nuances of early-life immune responses. Instead, an “immune layering” view^97^ considers how evolution has shaped immune development to address ever-changing environmental challenges across ages. The newborn must quickly distinguish sudden microbiome colonization from potential pathogens while also tolerating self and new dietary antigens. The mechanisms by which newborns navigate this complex environment are poorly understood and likely rely on immune components generated during fetal hematopoiesis^112^. Because newborns respond differently to infection and vaccination and are susceptible to B-cell-mediated diseases^113^, we need to understand the unique molecular programs that distinguish early-life human B-1 cell development and function. Although human B-1-like populations have been described in the fetal spleen^5^ and different groups have revealed B-1-like features in adults^56,114^, determining whether they emerge from the distinct developmental program we characterized here would provide further evidence critical for assigning a B-1-cell lineage identity. Thus, future investigations should link the fetal B-lymphopoiesis we describe here with the persistence, heterogeneity, and functions of these cells in postnatal life. Our studies here provide conserved gene signatures and stereotyped BCR sequences that may help track human B-1 populations in postnatal life and offer a new framework for assessing and targeting B-cell-mediated immune responses from childhood to adulthood.

## Methods

### Human Tissue acquisition and processing

We obtained human embryonic and fetal tissues from the Birth Defects Research Laboratory at the University of Washington with ethics board approval and maternal written consent. This study was performed in accordance with the ethical and legal guidelines of Emory University’s institutional review board. We determined donor age in days post-conception by foot length^115^. Donor sex was determined by PCR using SRY and amelogenin primers and confirmed by using sex-specific mRNA transcripts in the scRNA-seq data (Male: UTY, DDX3Y, USP9Y, ZFY; Female: XIST, TSIX). fL tissue was processed by mechanical dissociation using slides, washed, and incubated with 0.25% Collagenase I (07902, STEMCELL Technologies) in a 37 °C water bath for 5-10 min with intermittent vortexing, followed by incubation with cell dissociation buffer (13151014, Gibco) at room temperature for 5 min and filtration (70um filter). All experiments were performed in our custom-made RPMI-1640 media (def-RPMI-1640; deficient in biotin, L-glutamine, phenol red, riboflavin, and sodium bicarbonate)^116^ supplemented with 3% newborn calf serum (NBCS; 26010074, Gibco) with Benzonase (E1014, Sigma) unless otherwise noted. Fresh whole adult bone marrow (aBM) samples as well as cryopreserved mononuclear cells (MNC) and CD34^+^ cells were purchased from AllCells (Alameda, CA) and STEMCELL Technologies (Vancouver, Canada) or were collected from healthy adult donors through Emory University Hospitals under IRB00066294. Adult peripheral blood mononuclear cells (aPBMCs) were collected from healthy donors through the Children’s Clinical and Translational Discovery Core (Emory University). All subjects provided written informed consent before collection. We isolated mononuclear cells from fresh whole aBM and peripheral blood using the EasySep™ Direct Human PBMC Isolation Kit (19654, STEMCELL Technologies). For subsequent targeted enrichment of human fL, aBM, and aPBMC, we used EasySep Human B-Cell Enrichment Kit II without CD43 Depletion (19154 or 17963, STEMCELL Technologies) or EasySep Human CD34 Positive Selection Kit II (17856, STEMCELL Technologies). All samples were enriched except for one fL sample from 72 days post-conception. Table S1 includes additional metadata for the fL (7-20 wpc) and aBM (20-50 years old) samples.

### Fetal liver transplantation in humanized mice

Immunodeficient triple-knockout mice (TKO; B6.129S-*Rag2^tm1Fwa^ Cd47^tm1Fpl^ Il2rg^tm1Wjl^*/J) were obtained from the Jackson Laboratory (Strain #:025730; RRID: IMSR_JAX:025730). All mouse studies were approved by the Emory IACUC. Five days prior to transplant, TKO mice (4-5 weeks old) were preconditioned with anti-cKit-saporin^117–119^ (Biotin anti-mouse CD117; 105804, Biolegend) coupled to Streptavidin-ZAP (IT-27-1000, Advanced Targeting Systems) via retro-orbital intravenous injection, followed by sub-lethal irradiation (two doses of 2.25 Gy delivered four hours apart; RadSource RS2000 X-Ray Irradiator) 24 hours before transplant. CD34^+^ cells were enriched from cryopreserved fetal liver (142d donor) using the CD34^+^ positive bead selection kit (17856, STEMCELL Technologies EasySep Human CD34+ Selection Kit II). We administered 1.5 million CD34^+^ fL cells intravenously by retro-orbital injection, with co-transplant of 100,000 total mouse BM cells flushed from a TKO mouse femur immediately before transplant. Transplanted mice were kept on a Baytril (enrofloxacin) diet (MD300-050, Bio-Serv; Flemington, NJ) for four weeks post-transplant. Mouse blood was collected every 2-4 weeks by mandibular bleeds in K2 EDTA tubes (365974, VWR). Red blood cells were depleted using Ammonium-Chloride-Potassium (ACK) lysis buffer. At termination of the experiment, tissues from humanized mice were harvested. We flushed bone marrows and processed spleens by mechanical dissociation. We selected one mouse harvested at four months post-transplant for scRNA-seq analysis. Prior to single-cell encapsulation, murine CD45^+^ cells were depleted using anti-mouse CD45-PE antibody (103106, Biolegend) and MojoSort mouse anti-PE nanobeads (480080, Biolegend), followed by enrichment of human B cells by negative selection (17963, STEMCELL Technologies B-cell Enrichment Kit II without CD43 Depletion).

### Flow cytometry

Samples in single-cell suspension were stained with primary human antibodies and viability dyes (individual reagent information is provided in Table S10). Staining was routinely performed for 30 min on ice, protected from light, using defRPMI in 3% NBCS containing Brilliant Stain Buffer (566385, BD Biosciences). We ran samples fresh or fixed them in BD FACS lysis solution (1:6 dilution in sterile dH2O; 349202, BD Biosciences) for 30 min at room temperature. When intracellular staining was performed, the TrueNuclear kit (424401, Biolegend) was used according to manufacturer instructions. Flow cytometry data were acquired on a FACSymphony A3 or A5 using FACSDiva software (versions 9 and 10, BD Biosciences) unless otherwise noted. We analyzed data using FlowJo (v10.10.0).

### Single-cell library preparation and sequencing

We enriched samples for B cells as described above for fL, aBM, aPBMC and humanized mouse spleen. For ADT staining, up to 1 × 10^6^ cells per donor/sample were incubated with Fc block (422302, Biolegend) on ice for 10 min, followed by staining with a mix of oligo-conjugated (barcoded) antibodies (antibody details listed in Table S10) for 30 min on ice in RPMI-1640 media supplemented with 3% NBCS. After washing, cells were resuspended to 1200–1500 cells/uL and passed through a 20um nylon filter before loading onto a Chromium Controller (10X Genomics, Pleasanton, CA). We prepared gene expression, BCR, and ADT libraries according to the protocol for the respective 10X Genomics 5’ chemistry version. 5000-20000 cells were targeted for encapsulation using 5’v1 or 5’v2. We sequenced libraries on the Illumina NextSeq 550, NovaSeq 6000, or MiSeq platforms. Sample-specific sequencing details provided in Table S2.

### scRNA-seq Data Processing

We aligned reads using 10x Genomics Cell Ranger (v7.0.0 or v7.0.1) with the GRCh38 reference genome. We then processed the scRNA-seq data using a stagewise analysis pipeline centered on the Seurat v4.3.0 framework^120^. Raw 10X Genomics gene expression (GEX) and antibody-derived tag (ADT) data were loaded using Read10X(), and cell barcodes were annotated with sample identifiers. We normalized ADT data using centered log-ratio (CLR) transformation. To remove low-quality cells, samples were independently quality controlled using criteria such as percentage of mitochondrial content (percent.mt) and the number of UMI count for mRNA (nCount_RNA). RNA data was then normalized to 10,000 counts per cell using NormalizeData(). Detailed thresholds and filtering decisions by dataset are provided in the Table S2.

### Dimensionality reduction, clustering and integration

To generate the fL, aBM, and humanized mouse Uniform Manifold Approximation and Projection (UMAP), samples were first analyzed individually before integration. For initial per-sample analysis, we first identified highly variable genes (HVGs) using the variance-stabilizing transformation (VST) method and scaled them using ScaleData(). We performed Principal Component Analysis (PCA) on the scaled GEX data. We computed nearest-neighbor graphs and generated a two-dimensional UMAP visualization on the selected principal components (PCs), and identified cell clusters using the Louvain algorithm. We determined the number of HVGs (nHVGs), PCs, and clustering resolutions independently for each sample by checking the clustree v0.5.1 plot^121^(sample-specific parameters provided in Table S2). Clusters containing HS(P)Cs and B cells were manually identified based on the expression of canonical markers (HS(P)Cs: HLF, AVP; B cells: CD79A, CD19, MS4A1) and subsequently extracted as individual Seurat objects. Initial filtering removed non-B cell contaminants (e.g., TTR, KLRB1, LYZ-positive cells). HVGs were identified independently in each object (nfeatures = 900) and combined, excluding cell cycle-associated genes defined in “Seurat::cc.genes.updated.2019”.

To generate the integrated fL and aBM UMAPs, we used Harmony v0.1.1 batch correction^122^. We applied the correction via a custom RunHarmony.Seurat wrapper to the first 30 PCA dimensions to correct for sample-specific technical effects. We computed UMAP visualizations on the first 25 and 10 Harmony dimensions for fL and aBM, respectively. We performed Louvain clustering (resolution = 0.6 (10 fL); 0.9 (10 aBM)) on the Harmony dimensions to identify integrated clusters. The fL and aBM reference atlas mapping were generated using Symphony 0.1.1 via buildReferenceFromSeurat() after cell type annotation. The reference included saved PCA/Harmony embeddings, the UMAP model, and soft cluster assignments, enabling mapping of new fL query samples. For the humanized mouse spleen dataset, we generated three libraries from one humanized mouse spleen to increase cell count. We also performed batch correction across these three samples using Harmony, as described above.

### Reference mapping

We mapped yolk sac^77^ and embryonic liver^76^ progenitors to our fL object. We downloaded the Carnegie stage (CS)14 and CS15 liver data from GEO (GSE162950) and processed them the same way as the internal fL samples. The count matrix of the CS12 yolk sac was obtained directly from GEO (GSE268143). CD34^+^ cells were extracted and categorized as HSC (co-expression of HLF, MECOM, SPINK2, MLLT3, RUNX1, and HOXA9) or lymphoid progenitors (co-expression of SPINK2, JCHAIN, and LTB)^76^. scRNA-seq from the hPSC-derived B cells^93^ was similarly mapped to the fL object. We cleaned the dataset by removing progenitor cells expressing non-B cell lineage markers (CD3E, ITGAX, CD4, CDH11), and selected hPSC-derived B cells by cell-surface CD19 expression (ADT). We used Symphony embeddings to project the hPSC-derived B cells onto the established fL UMAP, as described above.

### Differential gene analysis

We used “FindMarkers” and “FindAllMarkers” from the Seurat package to identify differentially expressed genes (DEGs), and Wilcoxon rank-sum test with Bonferroni correction to determine statistical significance (adjusted p-value >0.05). DEG parameters and comparisons are listed in Tables S3-S9. To compare the mouse and human genetic programs, we extracted mouse hematopoietic progenitor cells (HPCs, Lineage^−^Sca1^+^c-kit^+^CD48^+^) from eL (E16.5) and aBM (8 weeks) from GEO (GSE128761)^71^. Genes with statistically significant differences between mouse eL and aBM were compared to those in humans. Dot plot visualizations of transcript expression were generated with dot size scaled by area and expression less than 5% of the cluster not shown (Seurat, v5.5.1, ggplot2, v4.0.3). Heatmap graphs depict average gene expression profiles (Seurat, v5.5.1) and were plotted using the ComplexHeatmap package (v2.26.1).

### Repertoire analysis

BCR-seq data was annotated using IMGT/HighV-QUEST (v3.6.1)^123^ to assign alleles and junctional features such as N-additions. We aligned reads with allele searching and insertion/deletion detection enabled. We processed IMGT outputs using our V(D)J analysis workflow^36^, available at https://github.com/Ghosn-Lab. We merged the processed Cell Ranger output with AIRR-formatted IMGT annotations by sequence_id (contig) to incorporate additional features, including junction N-addition and palindromic sequence details. Data cleaning was performed in R using dplyr (v1.1.4). We analyzed cells denoted high_confidence by Cell Ranger. Full-length and productive contigs annotated by IMGT were retained for analysis. We used the following criteria to select the primary heavy- and light-chain contigs per cell. When multiple productive heavy-chain contigs were present for a single cell barcode, we selected the contig with the highest unique molecular identifier (UMI) count. If multiple heavy-chain contigs displayed the exact same UMI, the cell was excluded. Light-chain contigs assignment was restricted to high-confidence productive IGK or IGL contigs from cells already containing a productive heavy chain. For cells containing both productive IGK and IGL contigs, the IGL contig was retained, consistent with presumed order of BCR rearrangement. Within the retained productive light-chain (IGK or IGL), we selected the contig with the highest UMI count and excluded cells with ties.

To compare Ig gene usage between datasets, we calculated the frequency of each observed VH, DH, and JH gene per dataset. The genes not detected in a dataset were assigned a frequency of zero. Finally, we calculated the Pearson correlation coefficient across the gene:frequency pairs between individual datasets to build a matrix of correlation values for visualization using the corr() function (stats package, v4.2.2). To evaluate CDRH3 properties, the Boman index was calculated using the CDRH3 amino acid sequence as input for the boman() function from the Peptides R package (v2.4.6).

### Humanized Mouse scRNA-seq analysis

To analyze B cells in the humanized mouse spleen, we selected B cells with productive heavy- and light-chain BCR expression. We further selected mature B cells based on CD79A expression and the absence of RAG1, RAG2, IGLL1, CD34, VPREB1, and DNTT. We then performed dimensionality reduction and clustering as described above. To assess fetal and adult gene expression, we performed differential expression analysis (DEGs) in mature B cells (cluster 8) between fL and aBM and obtained fL and aBM gene sets. To calculate the fetal score, we used the fL gene set to generate a UCELL score (AddModuleScore function in the UCELL package). We generated a score variation (fetal score 2) excluding SOX4 and used it when analyzing data subsetted on SOX4 expression. To classify antibody-secreting cells (ASCs), we also used a gene signature-based approach. We selected six positive marker genes associated with the plasma cell program and six negative marker genes associated with the B-cell program. A composite signature score was calculated per cell as the mean-normalized expression of the ASC genes (XBP1, CD38, CD27, PIM2, SDC1, PRDM1) minus that of the B-cell-associated genes (SPIB, BACH2, PAX5, FCER2, MARCH1, BANK1), using the log-normalized transcript data. For each cell, we summed total BCR UMI counts across all Ig productive contigs (“ighkl”). Cells with an ASC signature score greater than 0 and a total BCR UMI count (ighkl) exceeding 1,000 were considered ASCs. We also identified ASCs from the ten aBM samples and three additional PBMC donors using these criteria. To compare the fetal program in humanized mouse B cells with that of infant B cells, we downloaded human blood naïve B cells (GSE233321)^95^ from infants to adults (0-105 years old). scRNA-seq objects were converted from .h5ad files to Seurat .rds objects and normalized using Seurat’s LogNormalize method (scale factor = 10,000). We then calculated the UCELL fetal score for SOX4^+^ and SOX4^−^ infant naïve B cells.

### In vitro culture

Fetal liver and adult BM samples were enriched by CD34^+^ positive bead selection (17856, STEMCELL Technologies). Cells were then sorted (Cytek Aurora CS Cell Sorter, 100um nozzle; SpectroFloCS software v1.4.1) to obtain lineage-negative (Lin^−^) hematopoietic progenitors (CD45^+^CD34^+^CD3^−^CD14^−^CD16^−^CD56^−^CLEC4C^−^CD19^−^) and pro-B cells (CD34^+^CD19^+^CD24^+^). 4×10^4^ Lin^−^cells (48-well plate) or 1×10^4^ pro-B cells (96-well plate) were plated per well. After 16 hours, we determined a baseline live cell count by flow cytometry and initiated cytokine treatment. We treated cells with 10 ng/ml of the appropriate cytokine every three days along with half-media changes. Cells were split 1:4 (on day 6 and day 12) if wells were >80% confluent. After 15 days of cytokine treatment, plates were placed on ice and cells harvested using cold def-RPMI supplemented with 3% NBCS and Benzonase. Cells were stained according to the flow cytometry protocol above and run without fixation on the day of harvest.

### Antibody secretion quantification

We collected mouse blood from mandibular blood bleeds and centrifuged it at 300g for 30 min to separate plasma. We quantified human antibody concentration in humanized mouse plasma using our bead-based flow cytometry assay for antibody detection, FlowLITE, as previously described^124^. Briefly, yellow streptavidin-coated fluorescent beads (NC1863707, Spherotech) were coated with biotinylated anti-human isotype IgM, IgG, and IgA to saturation. 0.5 ul of plasma was incubated with coated beads, then revealed using PE-conjugated anti-human kappa and lambda antibodies. Data were acquired on a Cytek Aurora using SpectroFlo software (v4.0.3) and analyzed using FlowJo (v10.10.0).

### Statistical analysis

Statistical analysis was performed in R. Comparisons between groups were evaluated by two-sided Student’s t-test (base package) or linear mixed-effects model (lme4, v2.0.6) using donor as a random effect. Pairwise stage comparisons were performed using estimated marginal means (emmeans, v2.0.4) with multiplicity-adjusted p-values (Tukey method).

## Supporting information

Tables Index

## Acknowledgements

We thank the Emory Pediatrics/Winship Flow Cytometry Core (RRID: SCR_022324; NIH P30CA138292), the Emory Division of Animal Facilities, the Emory Children’s Clinical and Translational Discovery Core, and the following sequencing core facilities: Emory Integrated Genomics Core (EIGC, RRID:SCR_023529, NIH UL1TR002378), Emory Primate Center Genomics Core (NBRC, RRID:SCR_026418, NIH P51OD011132), Parker H. Petit Institute for Bioengineering, Bioscience at the Georgia Institute of Technology, Baylor College of Medicine Genomic and RNA profiling Core and Revvity Omics (PerkinElmer Genomics Inc.) for their support and technical assistance. We thank Dr. F. Eun-Hyung Lee, Sang N. Le, and Mindy R. Hernandez (Emory University) for providing select adult bone marrow samples. We thank the Ghosn lab members for their exploration of B-cell development biology; specifically, we thank David Cruz-Zarate for support with antibody measurements, Mackenzie (Landi) White for support with humanized mouse conditioning, Fathma Abdulkhader for tissue processing, and Justin Hahm for exploration of the B-cell receptor repertoire. We thank BDRL lab members, including Kimberly A. Aldinger, Dan Doherty, Ian G. Phelps, Jennifer C. Dempsey, Yasmeen Otaibi, and Lucinda A. Cort, for procurement of human tissues. We thank the Gordon Keller lab members (University of Toronto, Canada) for providing hPSC-derived cells. We used BioRender to generate select graphics.

This work was supported in part by the Gates Foundation (INV-044064, INV-071644, INV-073027, E.E.B.G.); the National Institutes of Health (NIH) National Institute of Allergy and Infectious Diseases (NIAID) R01AI182276 (E.E.B.G.) and R21AI167032 (E.E.B.G.); the National Institute of Child Health and Human Development (NICHD) R24HD000836 (I.A.G., Birth Defects Research Laboratory); the Emory Lowance Center for Human Immunology (E.E.B.G.); and the National Science Foundation (NSF) Graduate Research Fellowship Program (1937971/2439564, A.F.N.). Any opinions, conclusions, or recommendations expressed here are those of the authors and do not necessarily reflect the views of the NIH or NSF.

## Author contributions

This project was conceptualized by E.E.B.G. and A.F.N.; Wet-lab experiments were conducted by A.K. and A.F.N.; I.A.G. and BRDL researchers procured human tissue; Computational analysis was performed by K.K., J.Y., A.F.N., and P.C.; Data analysis were performed by A.F.N., K.K., J.Y., A.K., P.C., and E.E.B.G.; Formal analysis and visualizations were performed by A.F.N., K.K., J.Y.; A.F.N. and E.E.B.G. wrote the manuscript. All authors reviewed and accepted the manuscript.

---

**Figure S1.**
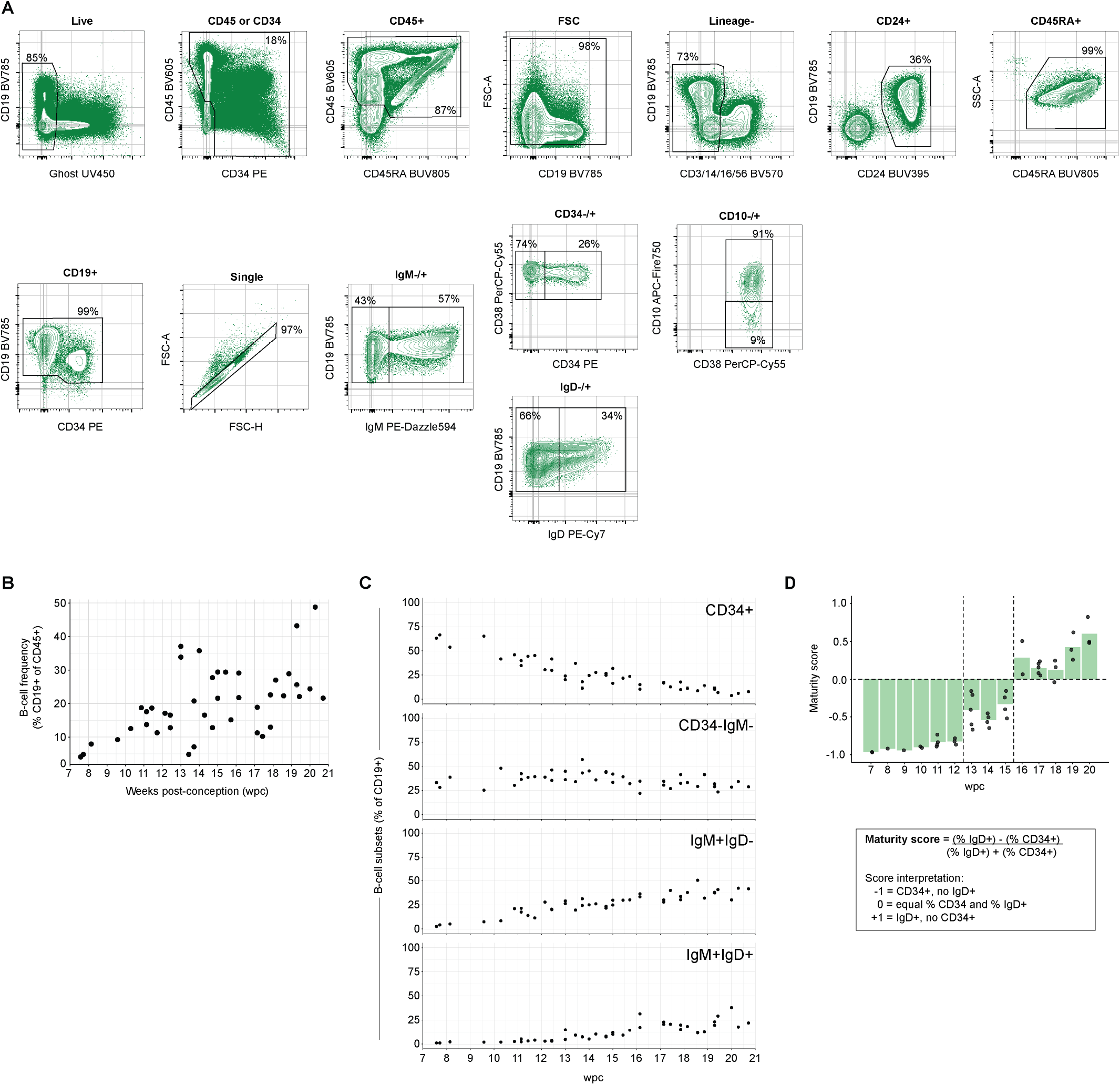
Developmental staging of human fetal liver B cells by flow cytometry. (A) Example gating strategy for developmental staging of fL CD19^+^ B cells. Live CD19^+^ B cells were distinguished by CD34, IgM and IgD expression. (B) B cell (CD19^+^) frequency in the fetal liver, determined as a percent of total CD45^+^ hematopoietic cells (n=42 fL donors). (C) B cell subsets across gestation. (D) Maturity score calculation assessing the percentage of early progenitors (CD34^+^) and late mature B cells (IgD^+^) within the fL B-cell population. Dashed lines denote gestational windows in which the ratio of CD34^+^ to IgD^+^ B cells changes.

**Figure S2.**
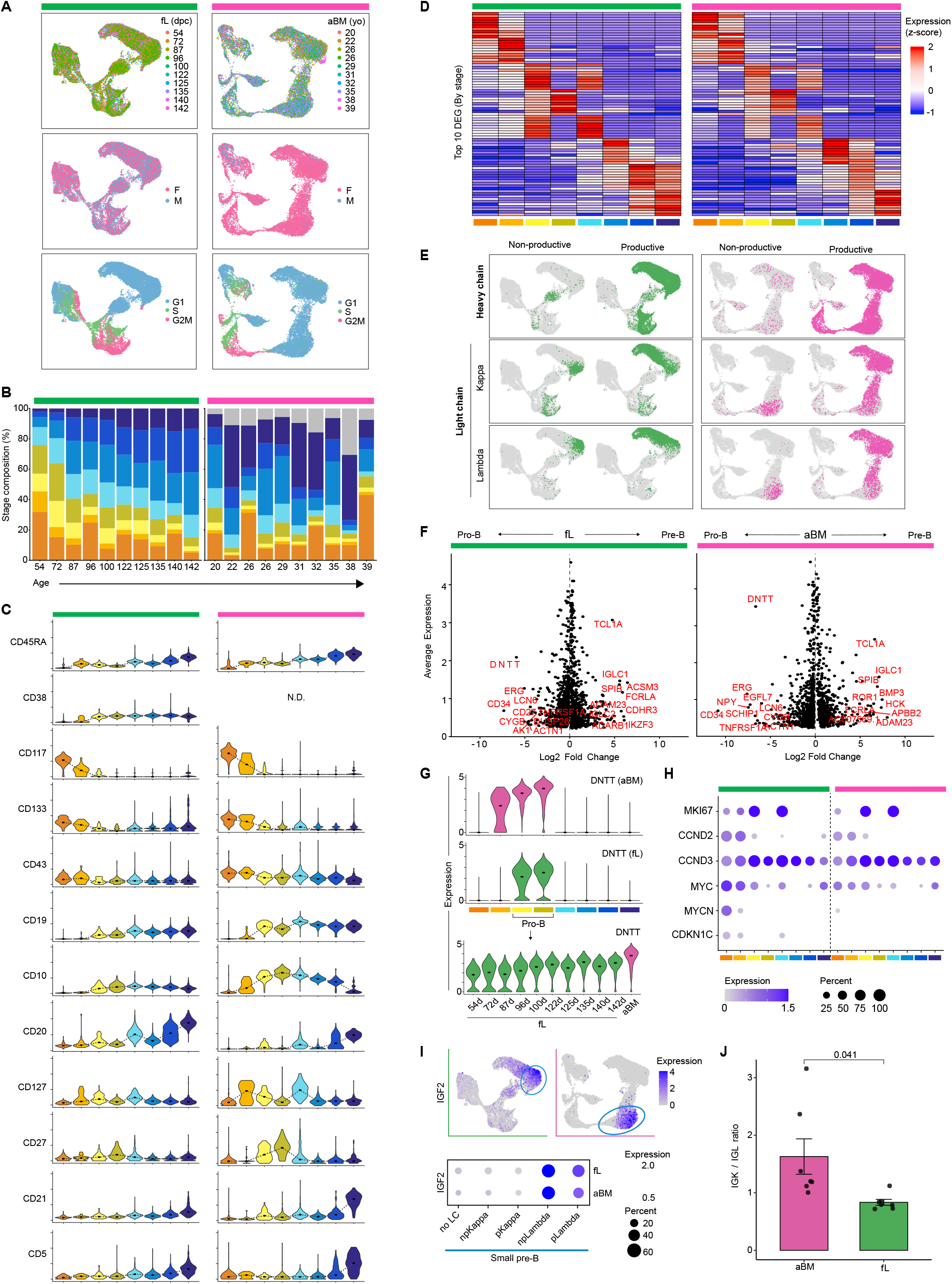
B-cell developmental staging determined by scRNA-seq. (A) Donor metadata displayed on fL (left) and aBM (right) UMAP embeddings. Characteristics (top to bottom): donor ID, donor sex, and cell cycle status. (B) B-cell stage composition by donor. Stages in Figure S2 (panels S2B, S2C, S2D, S2G, S2H) colored according to UMAP clusters in Figure 1C (left to right: HS(P)C, HPC, Early pro-B, Late pro-B, Large pre-B, Small pre-B, Immature B, Mature B). (C) Exemplary cell-surface expression (ADT) for fL (142d) and aBM (37yo) samples. Black dot denotes the median, with a dotted line indicating expression trends. (D) Top genes distinguishing B-cell stage in both fL and aBM. Overlapping stage-specific DEGs listed in **Table S3**. Top 10 genes shown, selected by average log fold change in fL. (E) VDJ status determined by BCR-seq projected on UMAP embeddings. (F) Pro-B (clusters 3 and 4) vs. pre-B (clusters 5 and 6) program in fL (left) and aBM (right). Volcano plot displaying DEG analysis (fold-change in expression between pro- and pre-B cells) and average gene expression (average log-normalized transcript in pro-B or pre-B cells, respectively). (G) DNTT expression in fL (green) and aBM (magenta) across stages of B-cell development (top). Expression in pro-B (clusters 3 and 4) displayed across age (bottom). (H) Dot plot displaying proliferation genes (percentages below 5% not shown). (I) IGF2 expression (log-normalized) within small pre-B subsets, determined according to progressive light chain rearrangement (kappa and lambda). (J) Ratio of kappa- to lambda-expressing cells within the HC^+^LC^+^ population. Donors with more than 500 HC^+^LC^+^ cells are included in the analysis.

**Figure S3.**
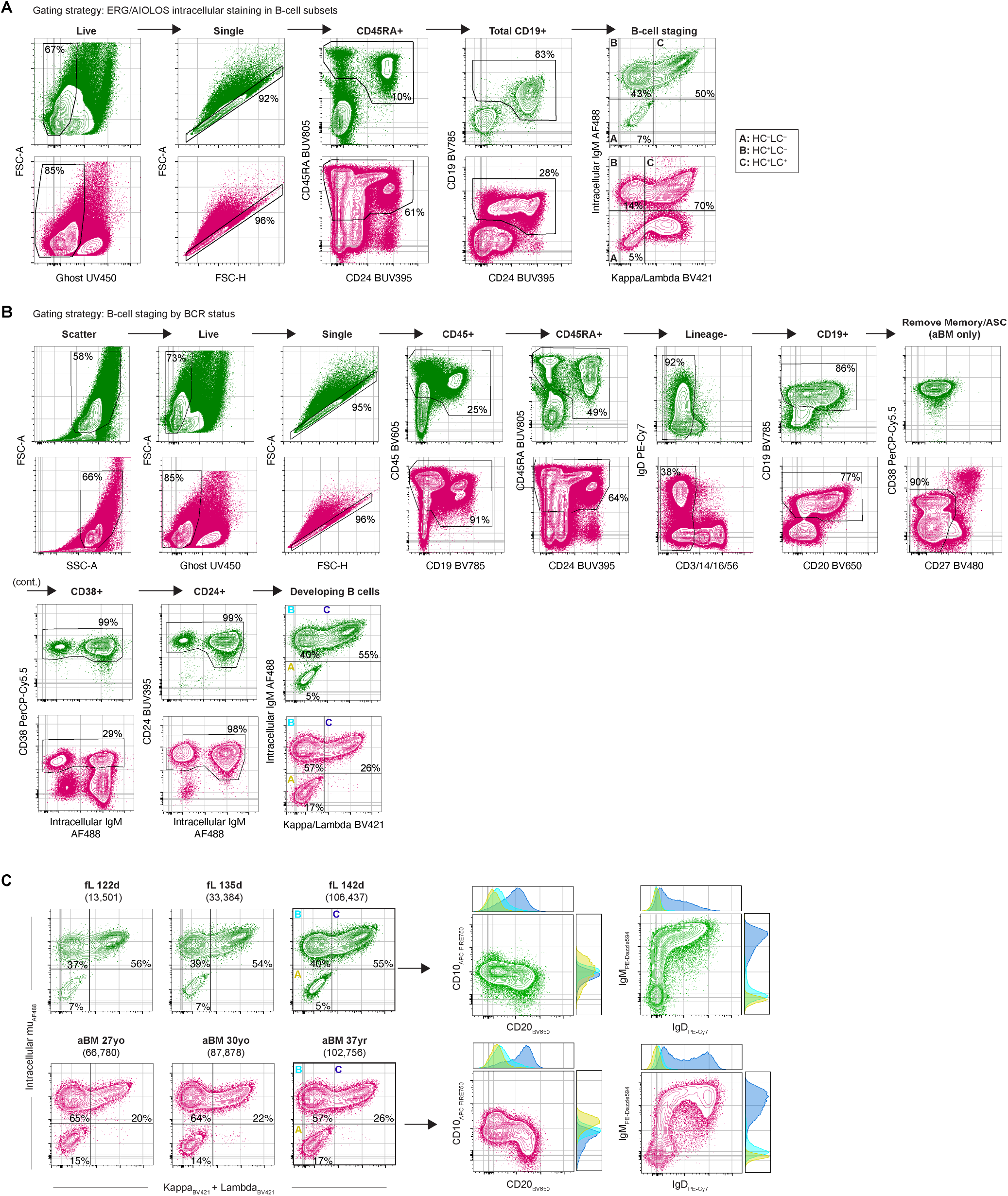
B-cell developmental staging according to B-cell receptor status by flow cytometry. (A) Representative gating strategy for CD19^+^ B cell selection for ERG and Aiolos staining analysis. Total fL cells (142d donor, green) and isolated mononuclear cells from aBM (37yo donor, magenta) are shown. (B) Representative gating strategy for staging of developing B cells using BCR status for fL (142d donor, green) and aBM (37yo donor, magenta). (C) Heavy chain (HC) and light chain (LC) subsets (populations A, B, C) in developing B cells (CD19^+^CD38^+^CD24^+^). CD10 and CD20, as well as IgM and IgD, reveal the developmental dynamics. Histograms display marker expression profiles in each B-cell subset: A: HC^−^LC^−^, B: HC^+^LC^−^, C: HL^+^LC^+^.

**Figure S4.**
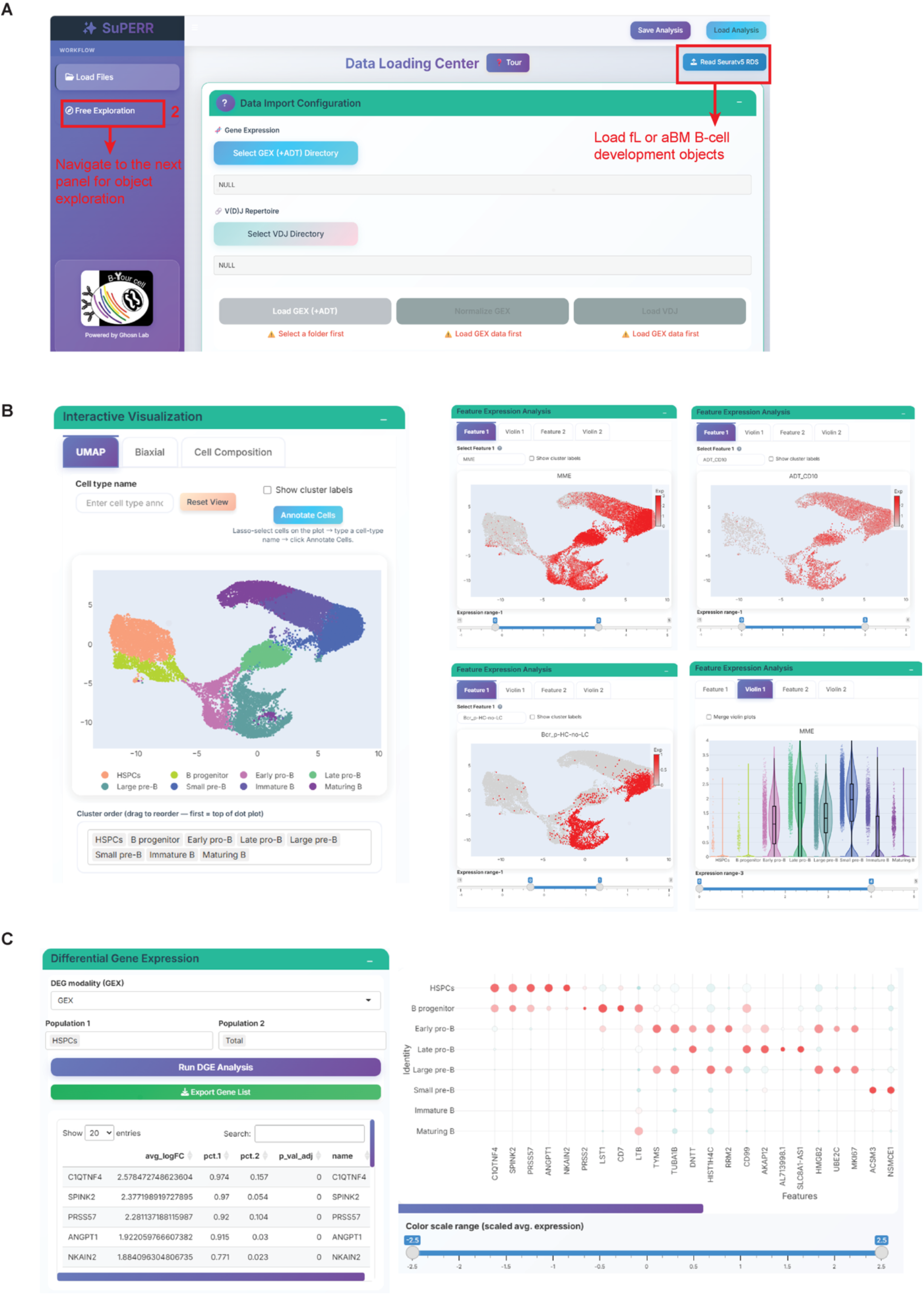
Interactive atlas of fL and aBM B-lymphopoiesis explored in the SuPERR R Shiny web application. (A) SuPERR is a user-friendly, open-source R Shiny application that runs locally and provides guided scRNA-seq data analysis and visualization. (B) SuPERR enables in-depth data exploration of B-cell development using GEX, ADT, and BCR datasets without requiring a coding background from users. (C) Flexible DGE analysis and dot plot visualization for the top 5 highly expressed genes per cell type.

**Figure S5.**
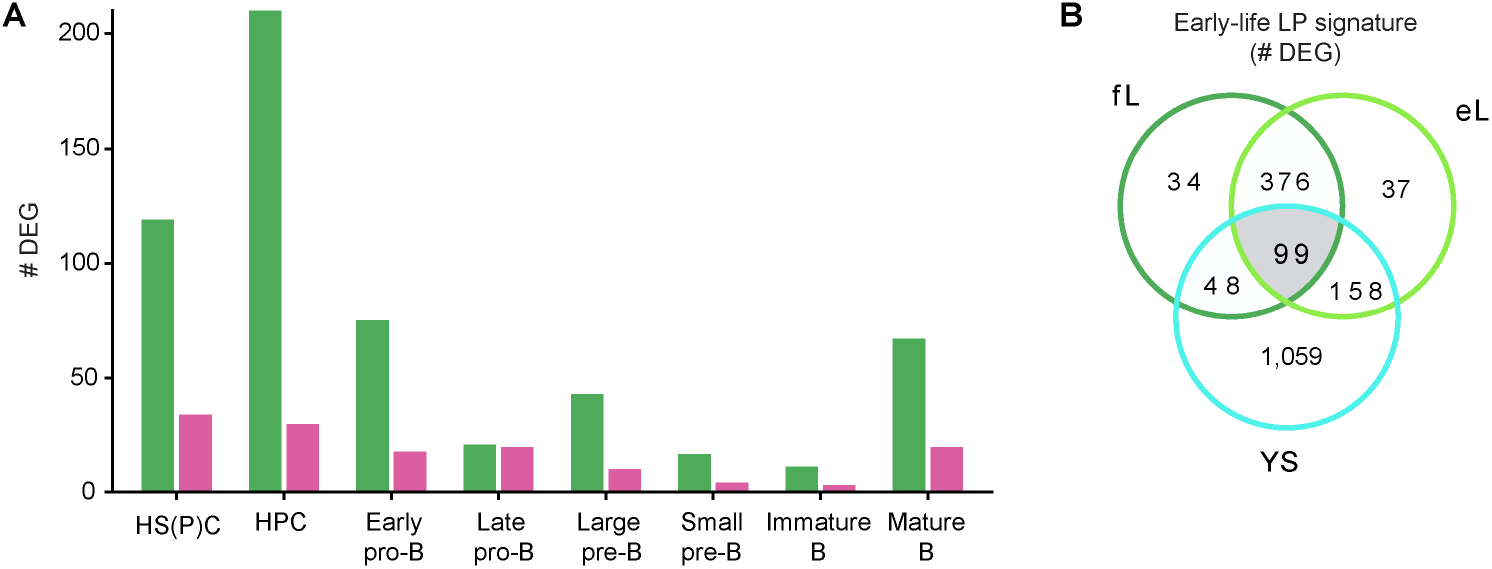
Early-life genetic program in lymphoid progenitors and developing B cells. (A) Tissue-specific differential gene expression (DEG) in fL (green) versus aBM (magenta) assessed across each stage of B-cell development. Number of DEGs by stage, determined by Wilcoxon rank-sum test using the Seurat FindMarkers function (DEG list included in **Table S5**). (B) Early-life signature determined by DEG analysis comparing to aBM lymphoid progenitors (LPs) (DEG list included in **Table S7**), resulting in 99 genes present in yolk sac (YS), embryonic liver (eL), and fL but not in aBM LPs.

**Figure S6.**
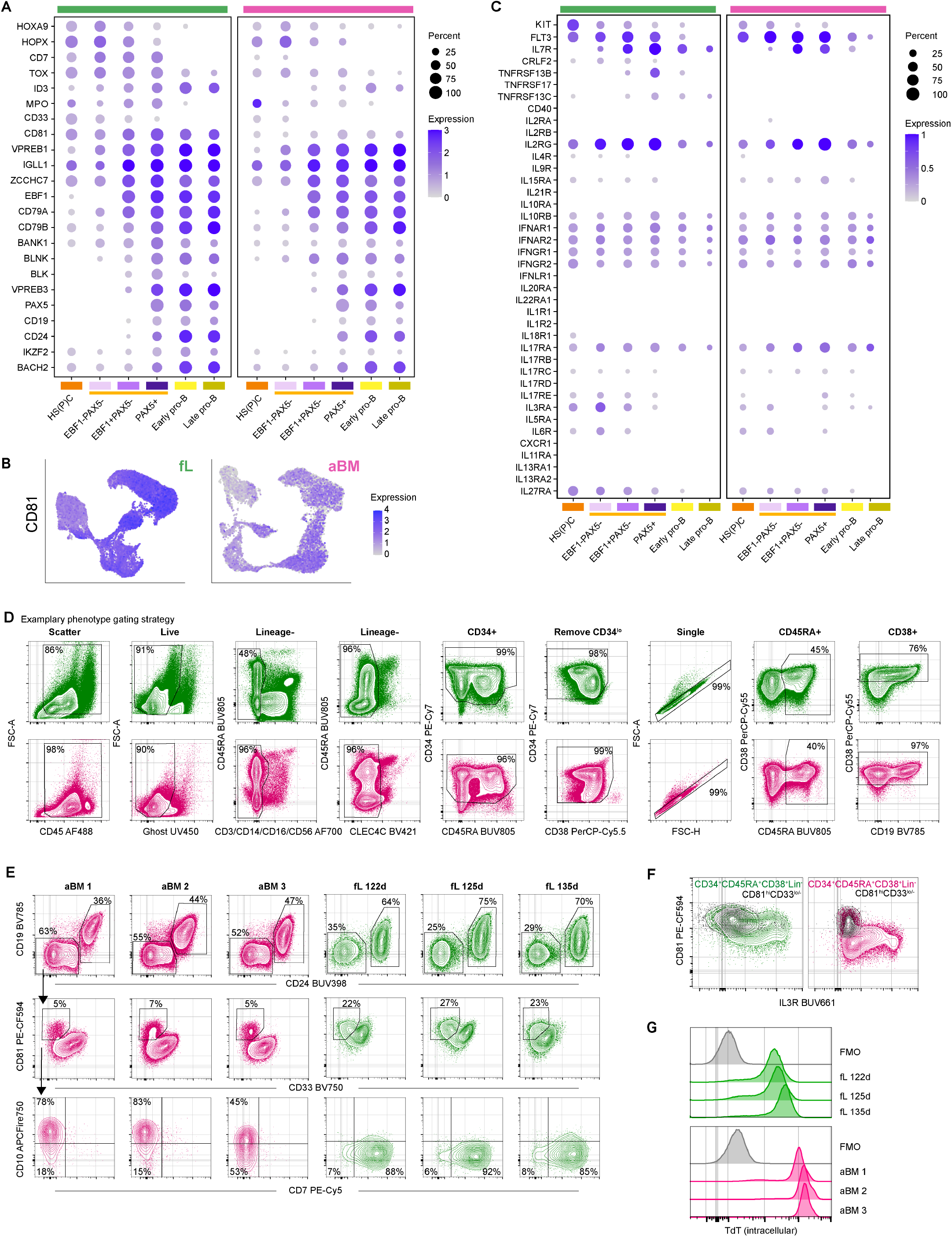
Characterization of early-life B-cell progenitors. (A) Expression (log-normalized transcript) of lineage commitment genes relative to EBF1 and PAX5. (B) CD81 expression (log-normalized transcript) is elevated in fL compared with aBM. (C) Cytokine receptor expression (log-normalized transcript). (D) Gating strategy for B-cell progenitors and pro-B cells from fL (125d donor, green) and aBM (43yo donor, magenta). CD34^+^ cells, after bead enrichment (STEMCELL Technologies), were stained and analyzed by flow cytometry (n=3 fL, 3 aBM). (E) B-progenitor and pro-B cell populations determined by CD24 and CD19 expression. Because fL has lower CD19 expression, CD24 expression helps define this population. CD19^−^progenitors can be further divided into myeloid CD33^hi^ and lymphoid CD81^hi^CD33^lo/–^ populations. (F) The B-lymphoid (CD81^hi^CD33^lo/-^) overlaid on the total lineage^−^ (Lin^−^) progenitor population (CD34^+^CD45^+^CD45RA^+^CD38^+^CD3^−^CD14^−^CD16^−^CD56^−^CLEC4C^−^CD19^−^). Despite broader and higher CD81 mRNA expression in fL, the lymphoid progenitors in both fL and aBM reach similar CD81^hi^ protein levels. (G) Intracellular staining of TdT in pro-B cells (CD19^+^CD24^+^) reveals lower TdT expression in fL compared to adults (n=3 fL, 3 aBM).

**Figure S7.**
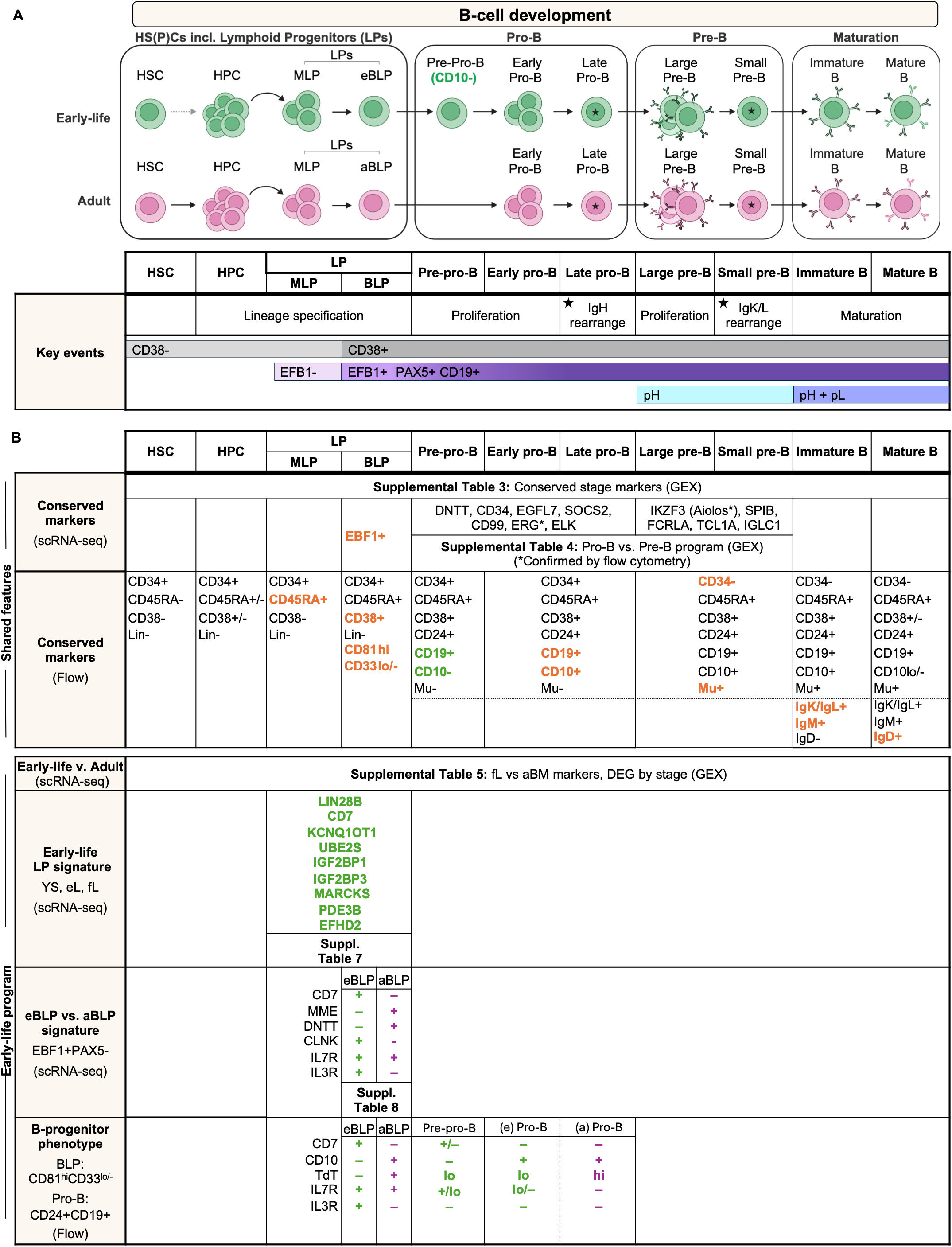
Revised maps of human B-cell development in early life and adults. **a,** Schematic of B-cell development stages displaying expression of key markers. The hematopoietic stem and progenitor cells (HS(P)Cs) include hematopoietic stem cells (HSCs) defined in scRNA-seq by their gene signature (HLF^+^MECOM^+^MLLT3^+^HOXA9^+^ RUNX1^+^SPINK2^+^)^76^. The first definitive HSC is generated in the embryonic aorta-gonad-mesonephros (AGM) around 4 wpc (∼32 days post-conception)^72^. Upon migration to the liver, fL HSCs undergo expansion and maturation^76^. It is unclear whether fL HSCs contribute to B-cell output or are primarily involved in maturation and dormancy, while multipotent progenitors support ongoing hematopoiesis^110,125^. Similarly, in mice^21,110,111^, studies show that embryonic-exclusive multipotent progenitors (eMPPs) develop independently of HSCs and are responsible for most fetal hematopoiesis, generating immune cells before definitive HSC output^21,110,111^. The human eMPP population would be contained within our HPC compartment and likely generates the eBLPs. We defined lymphoid progenitors (LPs) by scRNA-seq using their gene signature (SPINK2+LTB+JCHAIN+)^76^ and cell-surface protein expression (CD34+CD45RA+). The LP compartment contains the previously described multi-lymphoid progenitors (MLPs, CD38^−^)^81^ and early lymphoid progenitors (ELPs, CD38^+^)^1,9,10^. Our designation of BLPs was defined based on nascent Early B-cell Factor 1 (EBF1) expression, and thus may represent a subset within the ELPs, consistent with the observation that IL7R^+^ ELPs exhibit a strong bias to a B-cell fate^9,10^. EBF1 and Paired Box 5 (PAX5), followed by CD19 expression, mark progressive B-cell fate specification and commitment. Productive heavy chain (pH) and productive light chain (pL) are the components of the B-cell receptor (BCR) and are the result of successful V(D)J rearrangement. Predominant stages of active V(D)J rearrangement are indicated by black stars. **b,** Summary of B-cell development markers defined by scRNA-seq and flow cytometry.

**Figure S8.**
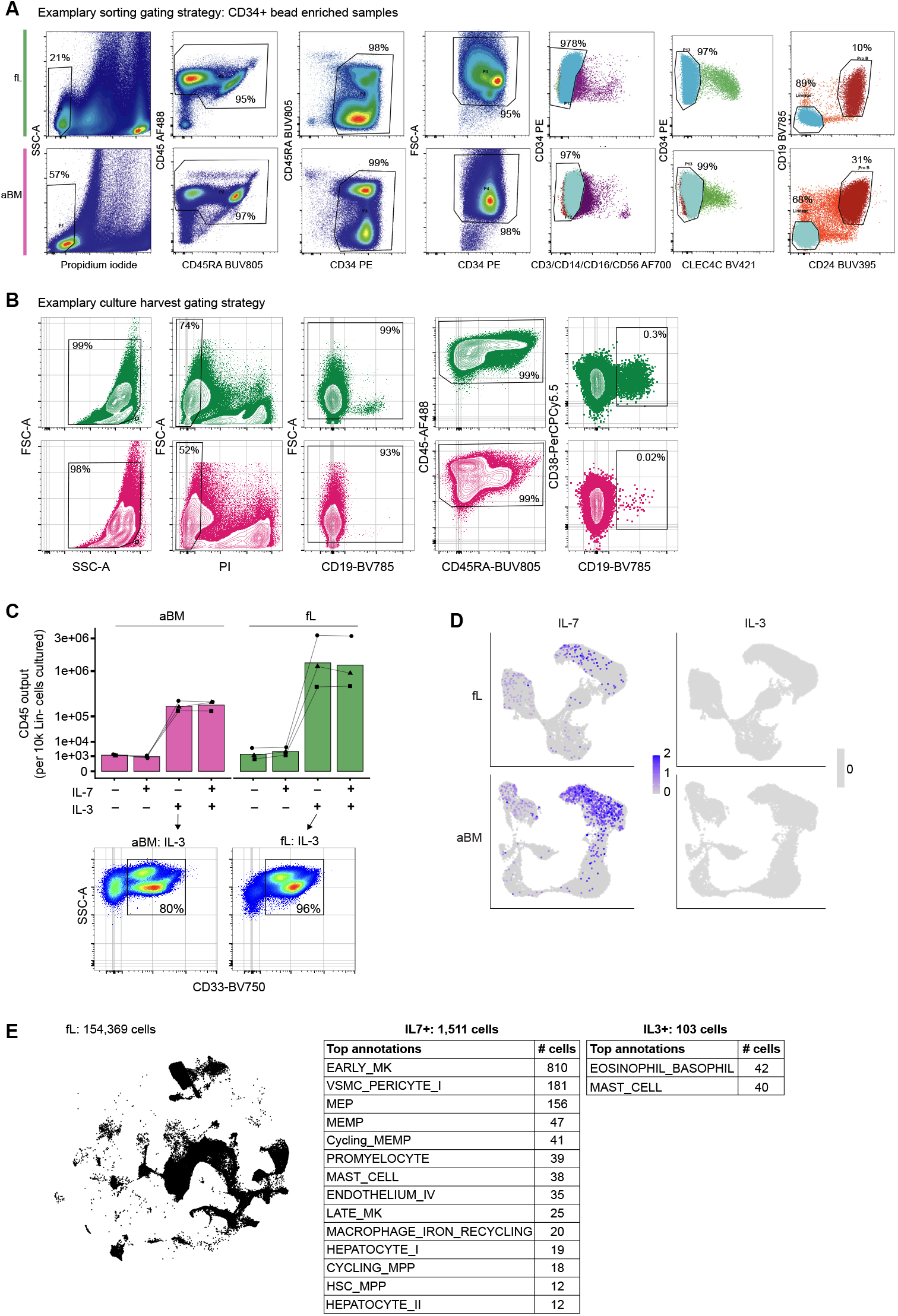
B-cell progenitor response to cytokines. (A) Representative sort strategy for the progenitor culture experiment (3 independent experiments, n=3 fL, 3 aBM). The first experiment included an additional removal of red blood cells via CD235a and doublets. (B) Representative gating strategy for CD19^+^ B cells generated after 15d in culture. (C) Total CD45^+^ cell output from lineage-negative progenitors after 15d in culture. IL-3-treated cultures show a large expansion of myeloid cells as indicated by CD33^+^ expression. Representative CD33^+^ expression in IL-3-treated fL and aBM cultures from Lin^−^ progenitors is shown. (D) IL-7 and IL-3 cytokine transcripts are displayed on the fL and aBM UMAPs. (E) Publicly available fetal liver (stroma and hematopoietic cells: 154,369 cells)^5^ was extracted for cytokine expression analysis. We excluded annotations for doublets, low-quality cells, high mitochondrial content, and erythrocytes. Cells with expression of IL-7 or IL-3 transcripts were selected. Cell annotations with 10 or more cells expressing cytokine transcripts were listed.

**Figure S9.**
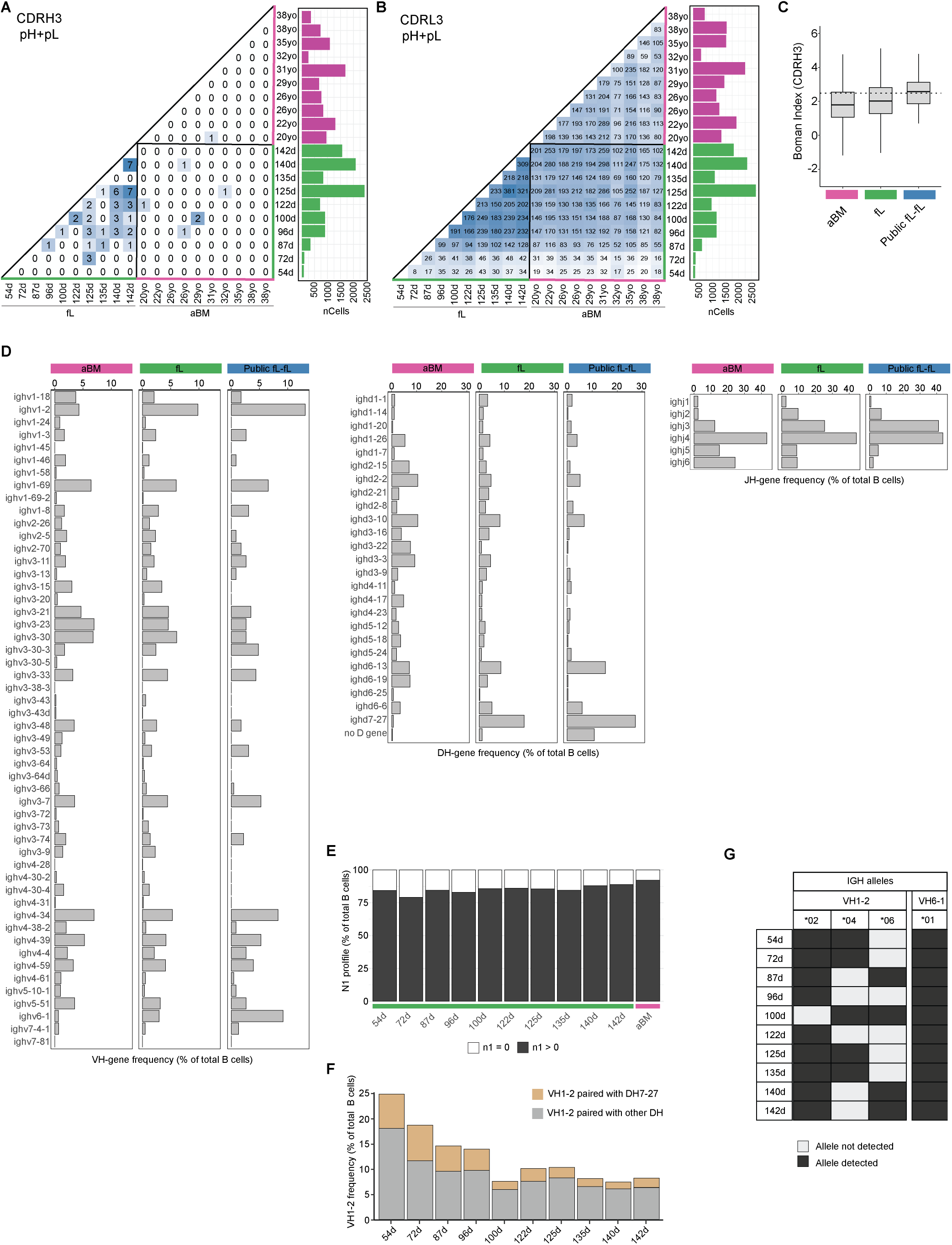
Features of the early-life B-cell receptor (BCR) repertoire. (A-B) Pairwise analysis of exact CDR3 amino acid (AA) sequence matches between donors in cells with both productive heavy and light chain (pH+pL) transcripts (9,879 fL cells and 11,125 aBM cells). (A) CDRH3. (B) CDRL3. (C) Boxplot displaying the Boman index for the CDRH3 sequences. The dotted line indicates the 2.48 kcal/mol cutoff, suggesting high protein-binding potential. (D) Heavy chain VH, DH, and JH gene usage. (E) N1-addition frequency across gestation. (F) Frequency of VH1-2 usage and pairing with DH7-27 across gestation. (G) Alleles detected in the ten fL donors for select VH genes (VH1-2, VH6-1).

**Figure S10.**
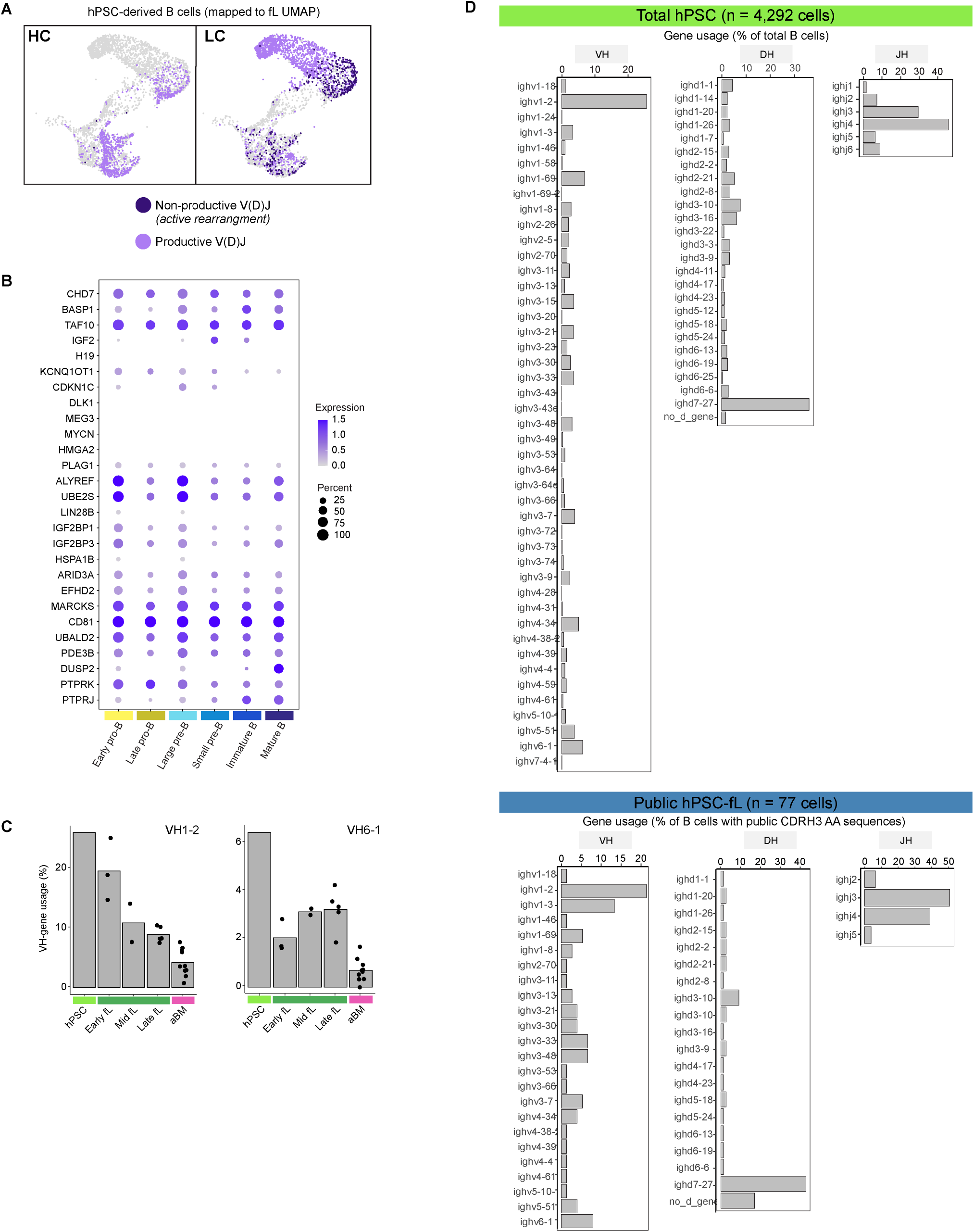
Characterization of hPSC-derived B cells. (A) Immunoglobulin gene**-**rearrangement status determined by BCR-seq, shown in a UMAP embedding of hPSC-derived B cells. (B) hPSC-derived B cells display a fetal-exclusive genetic program. Genes matching the fetal-exclusive program are shown in Figure 2A. (C) VH1-2 and VH6-1 gene usage across gestation. (D) Heavy chain gene usage (VH/DH/JH) in total hPSC-derived B cells (top) and in cells with public BCR (CDRH3 AA) matched between fL and hPSC (bottom). All cells with productive HC are included in panels **c-d**.

**Figure S11.**
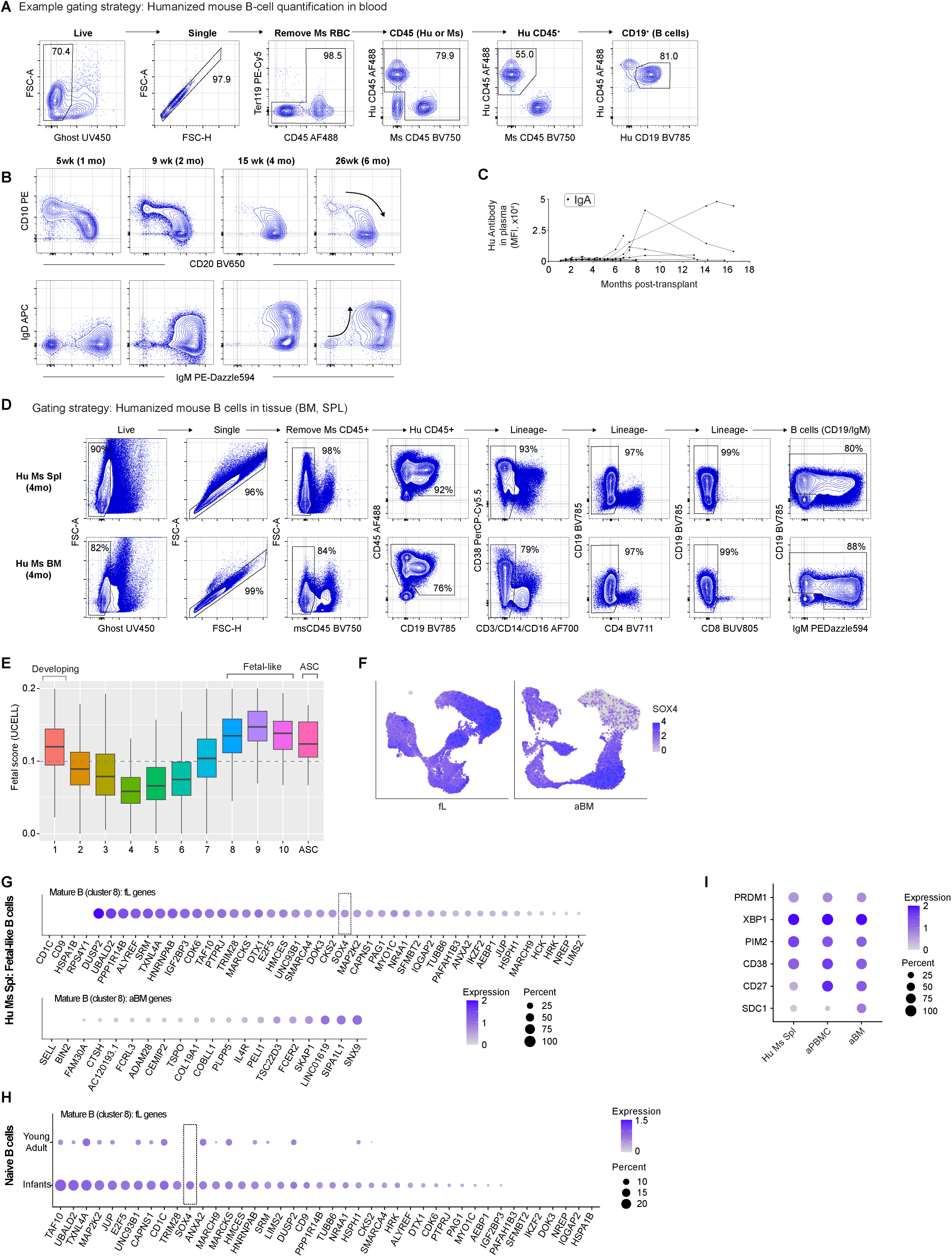
Development and differentiation of B cells in humanized mice. (A) Gating strategy to determine absolute counts of B cells in blood. We used counting beads to determine the CD19^+^ cell count per µL of blood. (B) Representative flow plots show B-cell development in blood over the first six months post-transplant. (C) IgA secretion in humanized mouse plasma measured by our bead-based flow cytometry assay, FlowLITE^124^. (D) Gating strategy to identify B cells and antibody-secreting cells (ASCs) in the humanized mouse spleen. (E) Fetal score (UCELL, derived from the fL cluster 8 gene list) in the mature Ig^+^ B cells from a humanized mouse spleen. Clusters with the highest fL score are annotated as fetal-like (clusters 8-10). (F) SOX4 expression (log-normalized transcript) along B-cell development in fL and aBM. (G) Gene expression in humanized mouse fetal-like cells. Gene list matching the differentially expressed genes in fL versus aBM mature cells (cluster 8). (H) Gene expression in infants (2months-2years, n=36) and young adults (18-40 years, n=24). (I) ASC gene signature in ASCs derived from the Hu Ms Spl (n=36 ASCs), human adult (a)PBMC (3 donors, n=70 ASCs), and human aBM (10 donors, n=548 ASCs).

