## Supplementary material for "Early-life human lymphopoiesis generates a distinct B-cell lineage": Tables Index

**Supplemental Tables Index**

| **Table** | **Title** | **Description** |
| --- | --- | --- |
| **Supplementary Table 1** | Donor information | Donor metadata |
| **Supplementary Table 2** | scRNA-sequencing datasets | scRNA-sequencing parameters |
| **Supplementary Table 3** | DEG: B-cell stage markers | DEG parameters: Min.pct=0.5, log2FC=1 (2x fold-change, in >50% of cluster), test=Wilcoxon  Selection: overlap between fL and aBM DEGs |
| **Supplementary Table 4** | DEG: Pro-B vs Pre-B | DEG parameters: Min.pct=0.25, log2FC=2 (4x fold-change, in >25% of cluster), test=Wilcoxon  Cell selection: Pro-B (Early pro-B & Late pro-B), Pre-B (Late pre-B & Small pre-B)  Selection: overlap between fL and aBM DEGs |
| **Supplementary Table 5** | DEG: Tissue exclusive GEX (fL vs aBM) | Log2FC=2, min.pct=0.25 |
| **Supplementary Table 6** | DEG: Mouse vs human HPCs | DEG parameters: tests=Wilcoxon  Human: HSPC fL vs aBM and HPC fL vs aBM: min_pct_1= 0.1, logfc_threshold=1, only.pos=T  Mouse: Ms eL HPC vs Ms aBM HPC: min_pct_1=0.1, logfc_threshold=1, only.pos=T  Selection: overlap between human fL HPCs and mouse eL DEGs |
| **Supplementary Table 7** | DEG: Early-life signature | YS/eL/fL vs. aBM  DEG paramaters: Log2FC=2, min.pct=0.25  Cells: lymphoid progenitors (SPINK2+LTB+JCHAIN+) in YS, eL and fL compared to aBM. |
| **Supplementary Table 8** | DEG: EBF1/PAX5 subsets | fL vs. aBM  DEG parameters: Log2FC=2, min.pct=0.25  Cell selection: HPC (cluster 2) split by EBF1/PAX5 in fL and aBM. |
| **Supplementary Table 9** | UCELL signature, DEG: Mature B (fL vs aBM) | DEG parameters: min.pct = 0.25, logfc.threshold = 2. test=Wilcoxon. fL signature was further refined by excluding genes expressed in more than 10% of aBM mature cells (pct.2<0.1). DEG matches Supplementary Table 3 mature B subset with an additional filtering. |
| **Supplementary Table 10** | Reagents details | Antibody information for ADT staining and flow cytometry. |
